# The 5α-reductase inhibitor finasteride reduces opioid self-administration

**DOI:** 10.1101/2020.09.15.291609

**Authors:** Gabriel D. Bosse, Roberto Cadeddu, Gabriele Floris, Ryan D. Farero, Eva Vigato, Suhjung J. Lee, Tejia Zhang, Nilesh W. Gaikwad, Kristen A. Keefe, Paul E.M. Philips, Marco Bortolato, Randall T. Peterson

## Abstract

Opioid use disorder (OUD) has become a leading cause of death in the US, yet current therapeutic strategies remain highly inadequate. To identify novel potential treatments for OUD, we screened a targeted selection of over 100 drugs, using a recently developed opioid self-administration assay in zebrafish. This paradigm showed that finasteride, a steroidogenesis inhibitor approved for the treatment of benign prostatic hyperplasia and androgenetic alopecia, reduced self-administration of multiple opioids without affecting locomotion or feeding behavior. These findings were confirmed in rats; furthermore, finasteride did not interfere with the antinociceptive effect of opioids in rat models of neuropathic pain. Steroidomic analyses of the brains of fish treated with finasteride revealed a significant increase in dehydroepiandrosterone sulfate (DHEAS). Treatment with precursors of DHEAS reduced opioid self-administration in zebrafish, in a fashion akin to the effects of finasteride. Our results highlight the importance of steroidogenic pathways as a rich source of therapeutic targets for OUD and point to the potential of finasteride as a new option for this disorder.

## Introduction

Over the last decade, the widespread abuse of prescription painkillers, such as oxycodone and hydrocodone, has led to a dramatic increase in the incidence of opioid use disorder (OUD) in North America ^1^. The extent of the crisis is such that opioids account for more than 60% of all drug overdoses in the United States, with an estimated 47 000 to 50 000 fatalities annually. Synthetic opioids, such as fentanyl, are the major drivers of opioid overdose^1,2^. Unfortunately, current therapeutic options for OUD are highly unsatisfactory. Existing treatments rely on replacement with long-acting opioids, such as methadone or buprenorphine^3^. While these options help patients cope with drug craving and manage withdrawal symptoms^3^, they are not ideal due to their intrinsic liability for abuse and dependence^4^. This background highlights the urgent need for novel, therapeutic options to reduce the risk and severity of OUD, particularly in patients requiring opioid treatment for chronic and neuropathic pain syndromes (for which effective alternatives to these painkillers are not always available).

Rodent models have been successfully used to model substance use disorders and study the circuitry and neurobiological changes involved in drug abuse^5,6^. However, the viability of these models to screen for novel therapeutic candidates is limited by their low throughput and high costs. In recent years, the zebrafish (*Danio rerio*) has emerged as a new alternative to study a wide range of complex behavioral and neuropsychiatric disorders, such as schizophrenia and depression^7–9^. Importantly, zebrafish have been shown to develop conditioned place preference and withdrawal symptoms after exposure to opioids and other drugs of abuse, such as cocaine and alcohol^10–14^. Additionally, adult zebrafish possess a complex central nervous system, including a blood-brain barrier, and considerable similarities with the mammalian homolog. Furthermore, this model is well suited for the rapid testing of candidate drugs, as compounds of interest can be dissolved directly into the water of the tank ^15–17^. Therefore, the zebrafish affords a unique opportunity to combine drug-discovery screening and substance abuse research.

Using a recently described paradigm to condition adult fish to self-administer the opioid hydrocodone^18^, we designed a behavior-based screen to identify novel compounds affecting opioid self-administration in zebrafish. We screened 110 unique molecules selected for their annotated activity against processes and pathways known to be involved in substance abuse disorders. From this screen, we identified the 5α-reductase (5αR) inhibitor finasteride (FIN)^19,20^ as one of the most effective compounds to reduce opioid self-administration. 5α-reductase catalyzes the rate-limiting step of the conversion of several ketosteroids, including progesterone and testosterone, into their neuroactive metabolites dihydroprogesterone and dihydrotestosterone (DHT). In turn, these steroids are converted into the neurosteroids allopregnanolone and 3α-androstanediol (3α-diol). (Figure 5A)^21–24^. Finasteride has been approved for over 25 years as a clinically approved treatment for benign prostatic hyperplasia and male-pattern baldness^25^. These effects reflect the suppression of DHT synthesis.

Our results showed that the same doses of finasteride that reduced opioid self-administration did not affect food-seeking behavior or overall locomotion. The effects on self-administration were confirmed in rats; furthermore, we identified the neurosteroid dehydroepiandrosterone sulfate (DHEAS) as a likely mediator of finasteride’s effects and found that finasteride does not reduce the painkilling properties of opioids. Using zebrafish, we thus uncovered a role for neuroactive steroids in the control of opioid self-administration that is conserved in mammals.

## Results

### Validation of the screening method

To screen for novel modulators of opioid self-administration, we utilized our newly developed assay to condition fish with the opioid hydrocodone. As previously described, small groups of adult zebrafish are conditioned to swim across an active platform to receive a dose of the drug^18^. Each visit results in the delivery of hydrocodone directly at the platform. Potential modulators of opioid self-administration are tested by training fish to self-administer hydrocodone for four days and treating with the compound of interest on the fifth day (Figure 1A). Before performing the small-molecule screen, we sought to validate the screening method by testing one of the only treatments used in the clinic, the slow-acting opioid methadone. Fish conditioned for four days to self-administer hydrocodone were treated with methadone (1 mg/L) for 60 minutes and transferred to the self-administration arena. Hydrocodone self-administration was significantly reduced in fish treated with methadone, when compared to control animals (Figure 1B), suggesting that the screening assay can identify molecules that reduce opioid self-administration.

**Figure 1.**
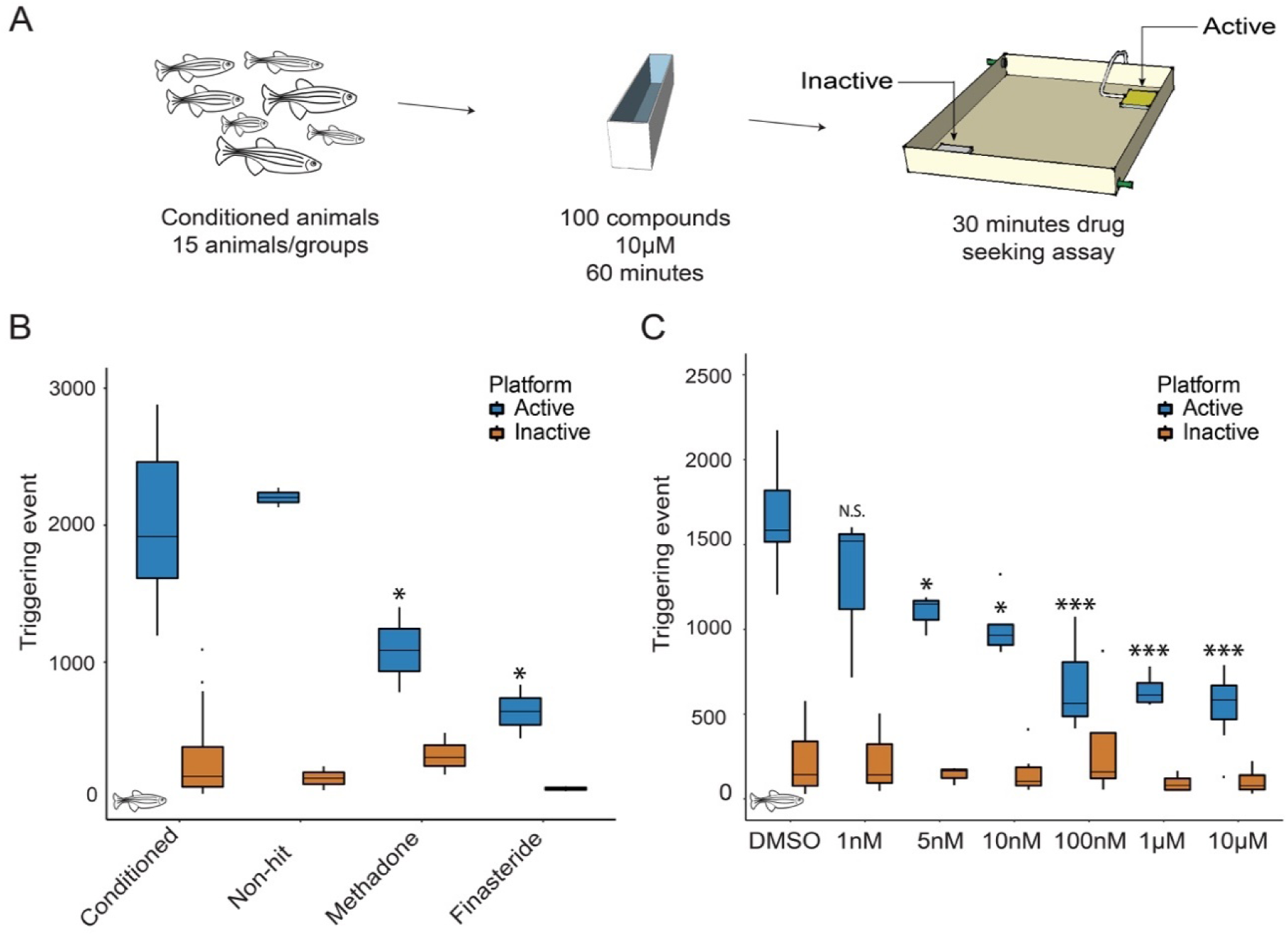
Small molecule screen for modulators of opioid self-administration. **A**. Conditioned animals were treated with the different molecules at 10µM for 60 minutes before assessing their opioid self-administration for 30 minutes. **B**. Known small molecules affect opioid self-administration. Treatment with Methadone (n=3), and Finasteride (n=2) significantly reduces the number of triggering events at the active platform. Conditioned fish n=56. No difference was observed at the inactive platform. *p*-value computed by Tukey HSD on one-way ANOVA, Inactive platform (F(5,64)=0.8711)p=0.50 and Active platform (F(5,64)=10.21)p=3.12E-07 compared to conditioned animals. **C**. Dose response experiment for Finasteride. Three doses, 100nM, 1µM, 10µM, reduce the number of triggering events below our threshold of 1000 activations. *p*-value computed by Tukey HSD on one-way ANOVA, Inactive platform (F(6,36)=1.06) p=0.40 and Active platform (F(6,36),24.60) p=2.31E-11, compared to the DMSO control. No significant difference detected for the inactive platform. DMSO n=16, 1nM n=3, 5nM n=5, 10nM n=6, 100nM n=5, 1µM n=4, 10µM n=7. * *p*-value <0.05, ** *p*-value <0.01, *** *p*-value <1E-5. Each n represents a group of 15 animals.

### Small molecule screening identifies modulators of opioid self-administration

We conducted a small-scale screen using a targeted collection of compounds with known activity on the regulation of neuronal and glial function, the immune system, or genes affected by substance use disorders. We hypothesized that focusing on this targeted collection would increase the probability of finding effective drugs within a smaller library of compounds. To test if the compounds reduce opioid self-administration, drug-conditioned animals were treated with 10 µM of each candidate compound 60 min before the 30-min self-administration session (Figure 1A). During each session, the number of triggering events for both the active and inactive platform was recorded and used as a readout of opioid self-administration. On average, control animals triggered the release of drug 1800 times per session; to reduce the number of false-positive hits, we tested each compound in duplicate. Furthermore, only compounds with less than 1000 triggering events for both duplicates were considered hits.

### Finasteride reduces opioid self-administration

After screening over 100 compounds, we identified the 5αR inhibitor finasteride as highly effective in reducing opioid self-administration. Incubation with a single dose of 10 µM of finasteride for 60 minutes was sufficient to reduce the number of triggering events at the active platform by 73% (Figure 1B).

To further validate that the inhibition of 5αR was responsible for the reduction in opioid self-administration, we tested a different 5αR inhibitor, dutasteride^26^. Similar to finasteride, dutasteride reduced the number of triggering events at the active platform (Figure S1). Although research on neuroactive steroids in zebrafish has been limited, the key steroidogenic enzymes are expressed in the adult brain, and the activity of 5αR has been detected in brain extracts^27–33^. Additionally, using a publicly available single-cell RNA seq library, we were able to detect 5αR transcripts in several different cell types in zebrafish brains^34^ (Figure S2). Taken together, these results suggest that the inhibition of the enzyme 5αR reduces opioid self-administration in zebrafish.

To further characterize the efficacy of finasteride, we performed a dose-response experiment. We used 10-fold dilutions to test finasteride concentrations from 10 µM to 1 nM. A significant difference was detected with concentrations as low as 5 nM (Figure 1C). This dose-response experiment supports the idea that finasteride is highly potent and has a large therapeutic window in zebrafish.

### Finasteride does not affect locomotion or food self-administration

To determine if the effect of finasteride was caused by sedation, we monitored the swimming speed of finasteride-treated animals. Finasteride did not reduce locomotion in comparison with DMSO (Figure S3). Importantly, we also did not observe any significant difference in the number of triggering events at the inactive platform between the finasteride-treated fish and control animals (Figure 1). These results suggest that finasteride specifically reduces the number of triggering events at the active platform without affecting overall locomotion.

Drugs of abuse are known to activate the reward pathway in the brain, which is a core contributor to motivated behaviors, such as feeding or reproduction^35^. We thus decided to test if finasteride was also affecting other motivated behaviors by measuring food self-administration in zebrafish. We first trained fish as previously reported for opioids, but used food instead of opioid as the reward. Food-conditioned animals were then treated with DMSO or finasteride for 60 minutes, and the number of triggering events at the active platform was measured. As opposed to hydrocodone self-administration, finasteride-treated animals exhibited no decrease in food-seeking (Figure 2A). These data suggest that finasteride does not affect all motivated behaviors and further validate that this drug does not impair locomotion.

**Figure 2.**
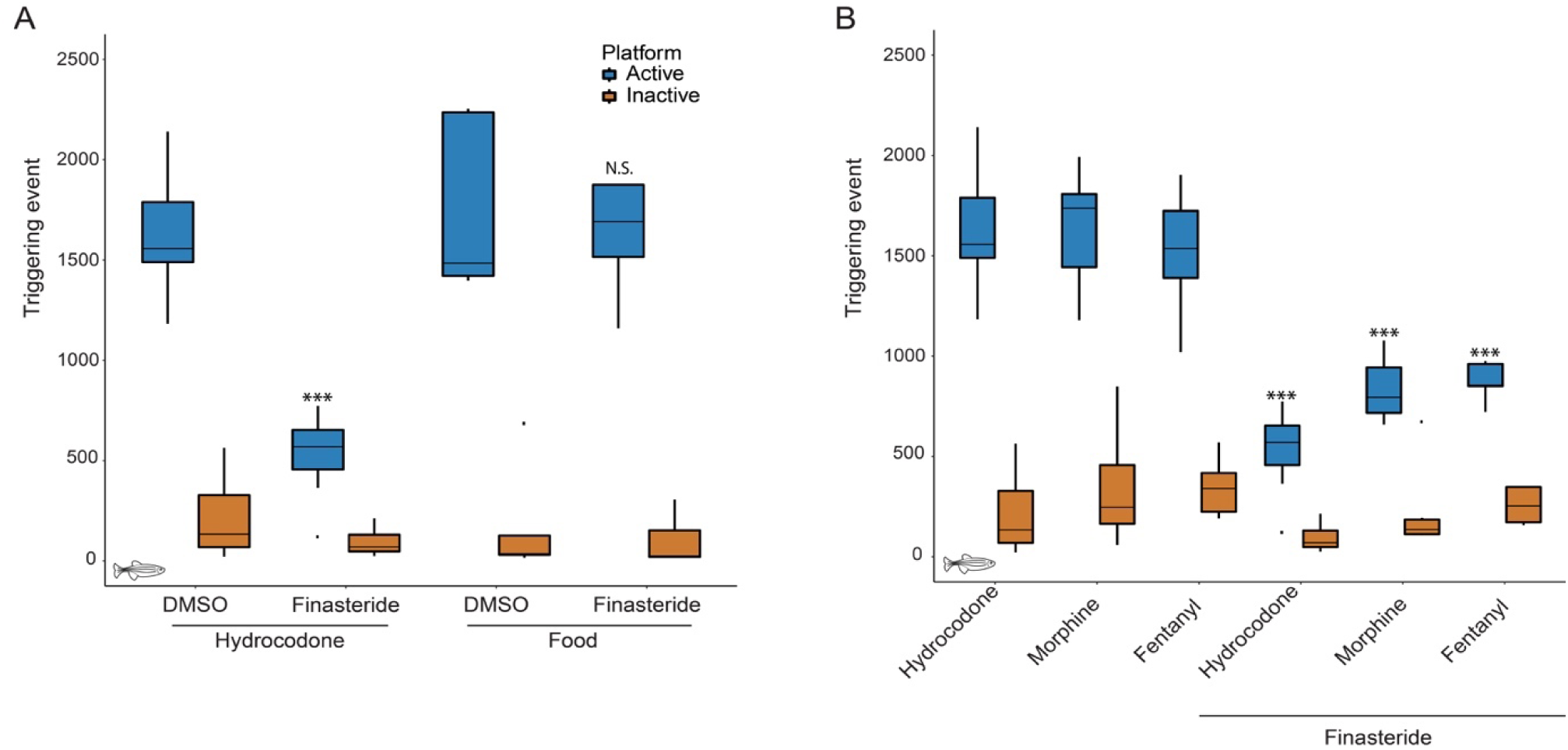
Finasteride effectively reduces the self-administration of different opioids, but does not affect all motivated behaviors. **A**. Finasteride reduces opioid self-administration without affecting food-seeking behavior. Finasteride significantly reduces the number of triggering events at the active platform compared to DMSO control for hydrocodone. No difference was detected for fish conditioned to self-administer food. *p*-value computed by Tukey HSD on one-way ANOVA, Inactive platform (F(3,29)=1.05)p=0.38 and Active platform (F(3,29)=19.37)p=4.32E-07, compared to respective DMSO control. Opioid trained animals: DMSO n=16, Finasteride n=7. Food seeking: DMSO n=5, Finasteride n=6. **B**. Finasteride affects opioid self-administration for animals conditioned with 3 different opioids. *p*-value computed by Tukey HSD on one-way ANOVA, Inactive platform (F(5,81)=1.57)p=0.18 and Active platform (F(5,81)=31.68)p=4.52E-13, compared to respective control. Hydrocodone control n=16, Hydrocodone+Finasteride n=7, Morphine control n=7, Morphine+Finasteride n=6, Fentanyl control n=7, Fentanyl+Finasteride n=5. No significant difference was detected for the inactive platform in any condition. * *p*-value <0.05, ** *p*-value<0.01, *** *p*-value <1E-5. Each n represents a group of 15 animals. Data for hydrocodone treatment alone reproduced from Figure 1C for comparison.

### Finasteride reduces self-administration of different opioids

It has been described that each class of opioids has different abuse potential^36,37^. Therefore, we decided to test the effect of finasteride on animals conditioned with the traditional opioid most commonly used in animal models, morphine, and one of the most potent and deadly opioids, the synthetic opioid fentanyl.

In order to test the effect of finasteride, we used the same conditioning protocol to train animals to self-administer morphine and fentanyl. As morphine and hydrocodone are closely related, we used the same dose of 6 mg/L. As for fentanyl, studies suggest that it is up to 50-100x more potent than morphine^38^, so a dose 50X lower was used, 0.12 mg/L to take into account its greater potency. We first confirmed that animals conditioned with either opioid had similar self-administration levels after 5 days of conditioning. There was no significant difference in the number of triggering events between fish trained with different opioids (Figure 2B). Therefore, this conditioning protocol is easily applicable to multiple classes of opioids. The conditioned animals were then treated with 10 µM finasteride for 60 minutes before performing a 30-minute self-administration session. As observed with hydrocodone-trained animals, there was a reduction in the number of visits at the active platform without affecting the inactive platform (Figure 2B). This result suggests that finasteride reduces opioid self-administration behavior regardless of the opioid used during the conditioning phase.

### Finasteride reduces opioid self-administration in mammals

The zebrafish model suggests that finasteride strongly reduces opioid consumption. To confirm this effect in mammals, we tested finasteride in a rat model of hydrocodone self-administration. Male rats were first conditioned to press a lever to receive an intravenous infusion of hydrocodone (0.016-0.128 mg/kg/160µL). Operant conditioning consisted of three stages of fixed-ratio (FR) reinforcement schedules: FR1, FR2, and FR5 (i.e., FR1=each lever press resulted in a hydrocodone infusion). Animals progressed to the next stage of conditioning after reaching criterion of >70% lever presses being on the lever on the previous schedule (Figure 3A). Importantly, to mimic the zebrafish conditions, rats had access to the drug for 60 minutes per day. To find the optimal concentration for hydrocodone conditioning, we tested a range of doses from 0.016 mg/kg to 0.128 mg/kg. We found that animals conditioned with 0.032 mg/kg and 0.064 mg/kg had the highest number of lever presses after 31 days of training (Figure 3B). We then treated animals conditioned with each concentration of hydrocodone by intraperitoneal injection of finasteride (50 mg/kg, i.p.) or vehicle before a self-administration session. That dose of finasteride was sufficient to significantly reduced the number of active lever presses at both 0.032 and 0.064 mg/kg/infusion of hydrocodone (Figure 3B).

**Figure 3.**
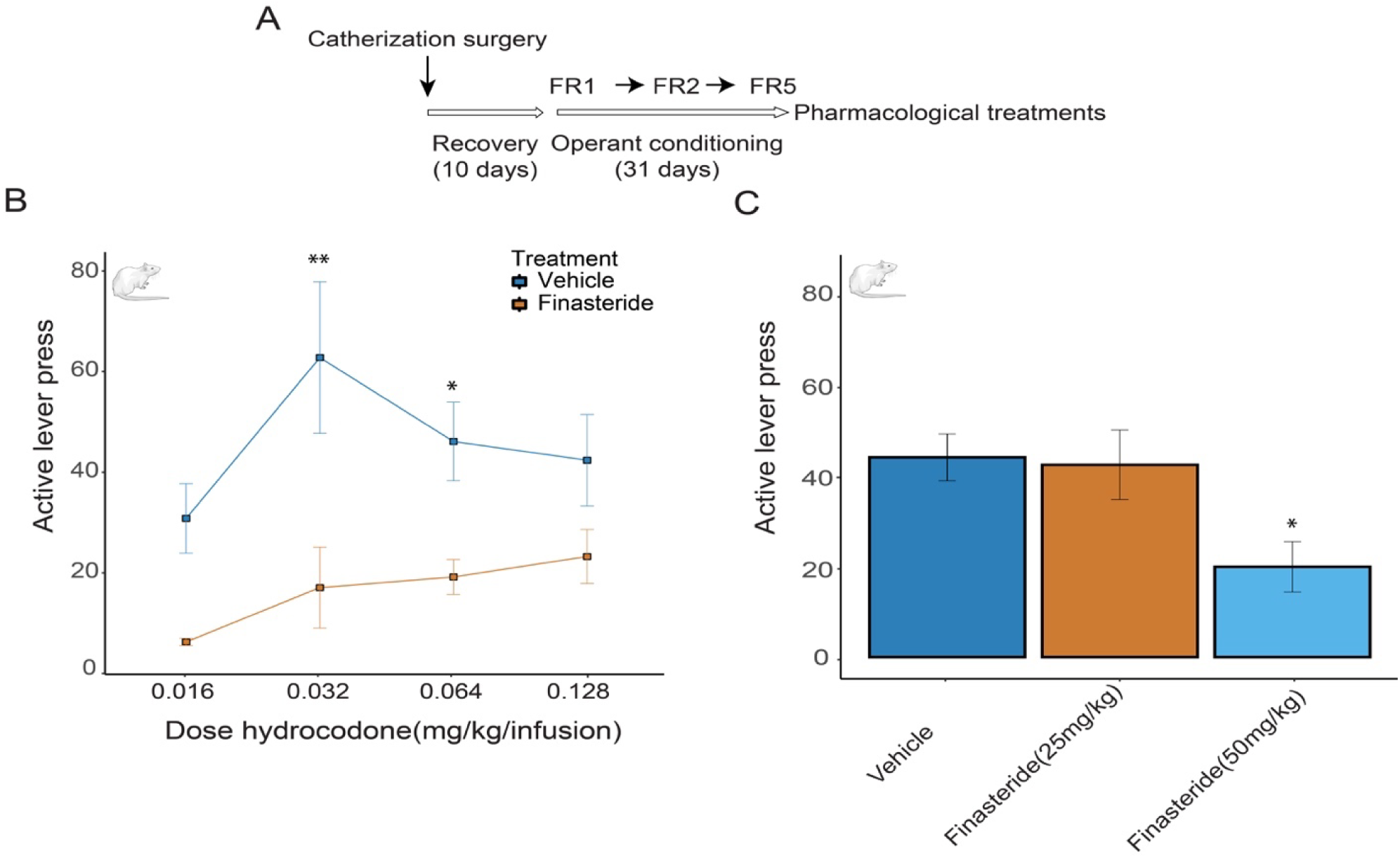
Opioid self-administration is reduced by finasteride treatment in rats. **A**. Rats were conditioned to self-administered hydrocodone and after establishing robust FR5, they were treated with either vehicle or finasteride. **B**. Active lever presses for animals trained with different concentrations of hydrocodone. Treatment with finasteride (50mg/kg) reduces opioid self-administration of hydrocodone for animals conditioned with 0.032 and 0.064 mg/kg. *p*-value corrected for multiple comparison two-way Anova (F(1,38)=30.5)p=0.0001) n=6 per condition. *p* value compared vs vehicle-treated animals \**p*-value < 0.05, ** *p*-value<0.01. **C**. IP injection with 50 mg/kg, but not 25 mg/kg finasteride reduces the number of active lever presses for *0*.*064 mg/kg hydrocodone*. n=8 animals. *p*-value corrected for multiple comparisons on Anova (F(2,21)=4.69)p=0.02. Error bars represent the mean +/- s.e.m.

To determine if finasteride was effective at different doses, we treated animals conditioned with 0.064 mg/kg of hydrocodone with 25 mg/kg or 50 mg/kg of finasteride. We confirmed that injection of 50 mg/kg significantly reduced the number of active lever press, but there was no difference in animals treated with the lower dose (Figure 3C). Importantly, locomotion or inactive lever presses were not affected by the treatment with finasteride (Figure S4 and S5).

To further validate that finasteride can reduce opioid intake in mammals, we also tested the effect of finasteride on the potent synthetic opioid fentanyl. Adult rats were trained to perform a nose-poke to trigger the release of a drop of fentanyl drinking solution (0.02 mg/kg/delivery) in a liquid magazine tray. We trained both male and female animals over 15 sessions of 60 minutes. As the training progressed, we observed an increase in active nose-pokes per session, and animals rapidly learned to discriminate between the active and inactive nose-poke holes (Figure 4A). Following each self-administration session, the mg/kg of fentanyl consumed was calculated for each animal by subtracting the amount of fentanyl left in the magazine tray from the total amount of drug delivered. As observed for the number of active nose-pokes, we also measured an increase in ingested fentanyl over time (Figure S6A).

**Figure 4.**
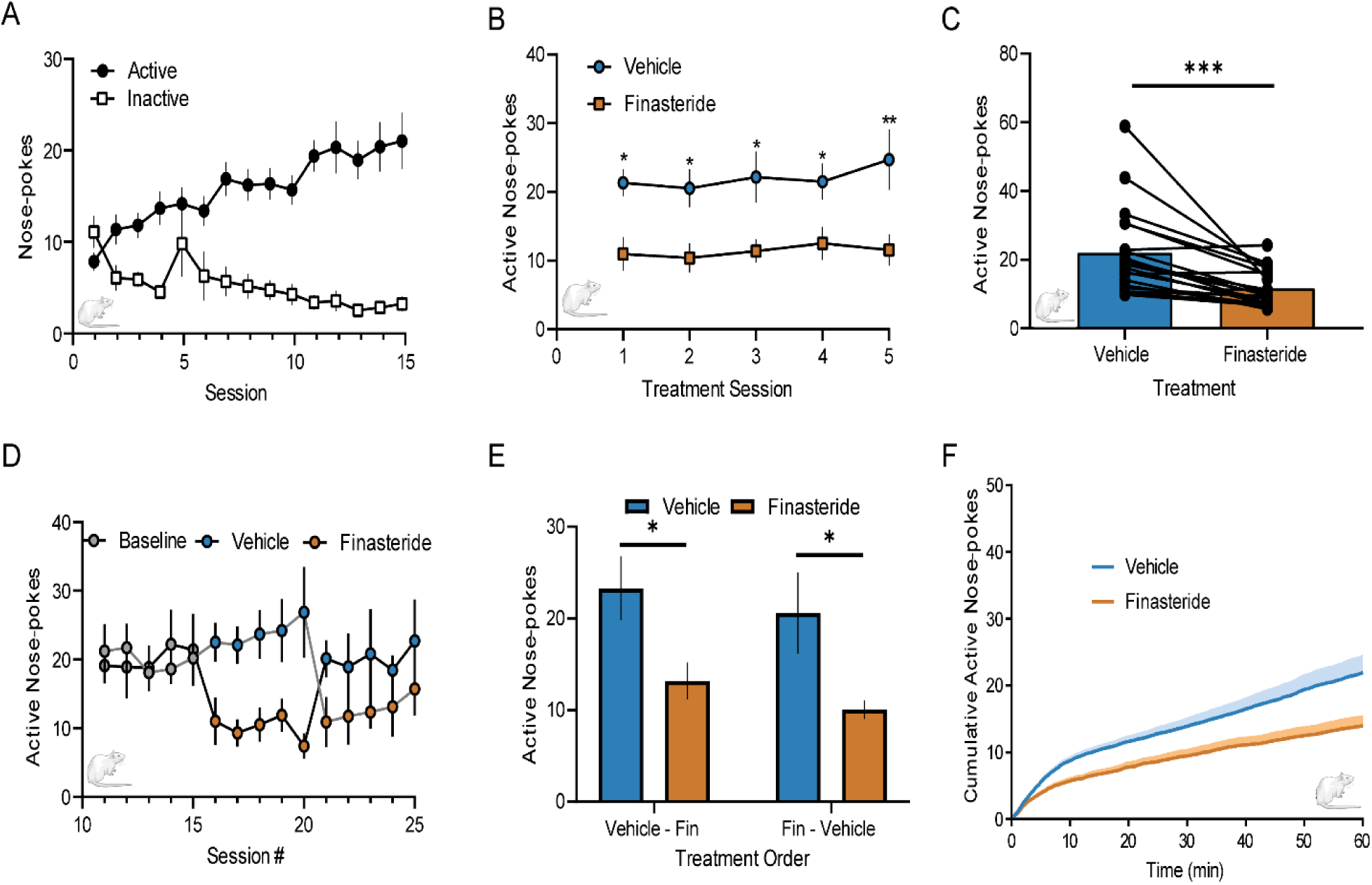
Finasteride decreases operant responding for oral fentanyl self-administration. **A**. Operant nose-poke responses at the active (blue circles) and inactive (orange squares) nose-poke ports during baseline fentanyl self-administration sessions (n=20 rats). **B**. Daily finasteride injections (IP, 50mg/kg) (orange squares) decreased active nose-pokes compared to vehicle injections (blue circles). A main effect of drug treatment was observed, Sidak post-hoc analysis was performed for multiple comparisons on mixed-effect model (F(1,19)=20.77)p=0.0002. **C**. Average operant responses during vehicle (blue) and finasteride (orange) treatment for each individual animal. A paired t-test revealed finasteride significantly decreased animals’ active responding for fentanyl delivery (t(19)=4.481)p=0.0003. **D & E**. Animals received daily injections of finasteride during sessions 16-20 (n=10 rats, D: black line, E: Finasteride-Vehicle) or during sessions 21-25 (n=10 rats, D: gray line, E: Vehicle - Fin). **D**. Baseline refers to animals responding during sessions 10-15. **E**. Order of treatment had no interaction with the effect of the treatment (F(1,18)=0.007)p=0.9321. A main-effect of treatment was observed (F(1,18)=19.03)p=0.0004. Sidak analysis was performed for multiple comparisons on two-way ANOVA. **F**. An average cumulative plot of active nose-poke responses during the vehicle-(blue) or finasteride-(orange) treated sessions (n=20). \**p*-value < 0.05, ** *p*-value<0.01, \*\*\**p*-value<0.001. Error bars represent the mean +/- s.e.m.

**Figure 5.**
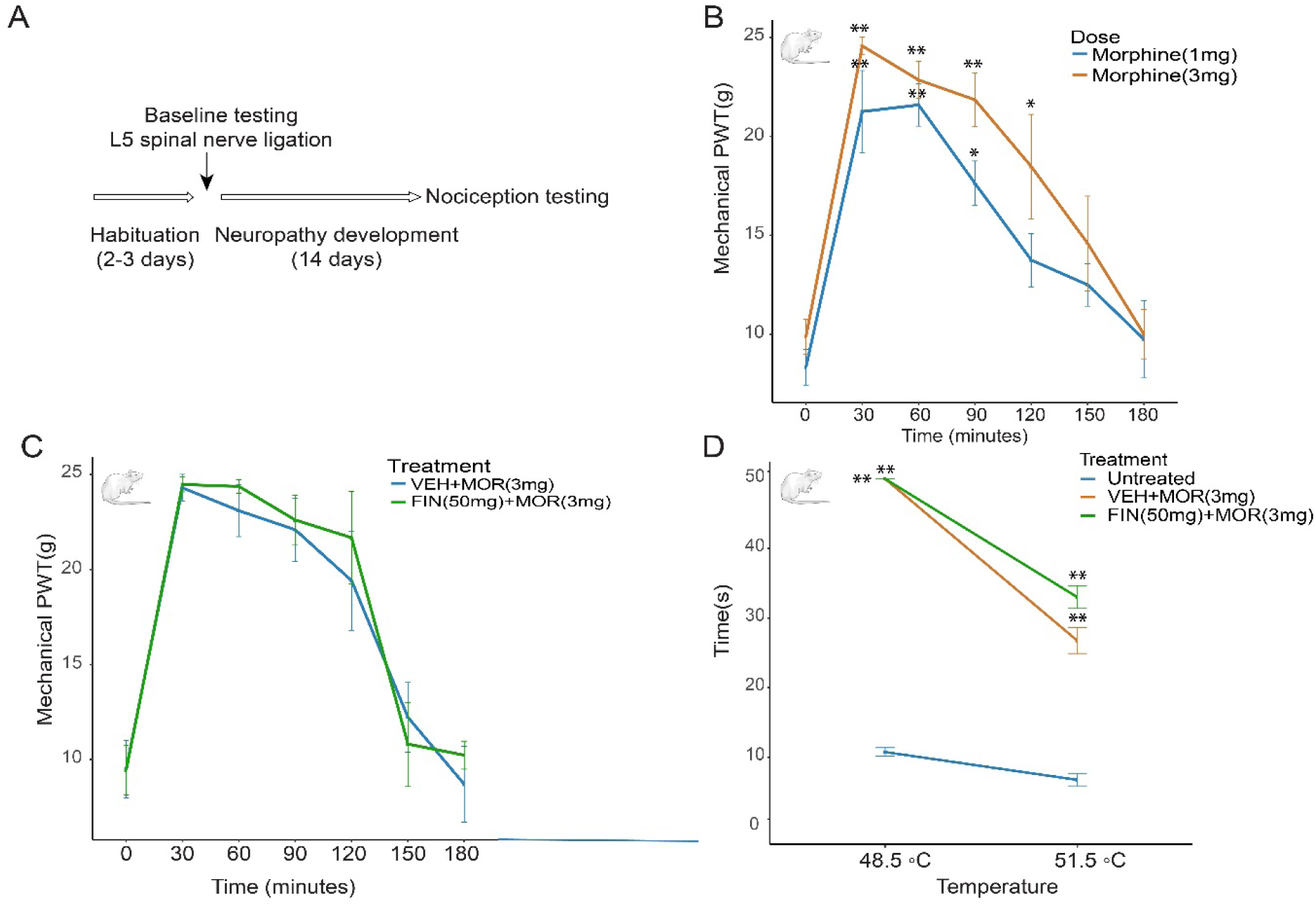
Finasteride does not affect the antinociceptive effect of opioids in a neuropathic pain model. **A**. After a 2-3 day habituation period, animals were assessed for baseline nociception tolerance followed by surgical L5 spinal nerve ligation. Neuropathy was established for 14 days before animals were tested for nociception with the different treatments. **B**. Paw withdrawal thresholds (PWT) to the Randall-Selitto test. Pain tolerance was measured over 180 minutes after treatment with different doses of morphine. Both doses of morphine significantly increased PWT compared to animals tested immediately after injection. Two-way ANOVA significant effect of Time (F(6,72)=31.09) p<0.0001. Morphine 1mg/kg, n=6, Morphine 3mg/kg, n=8 per condition. **C**. Co-treatment with finasteride (50mg/kg) did not block the antinociceptive effect of morphine (3mg/kg). Two-way ANOVA significant effect of Time (F(6,48)=45.31)p<0.0001, but no significant effect of treatment (F(1,8)=0.1875)p=0.6764 and no effect of interaction (F(6,48)=0.3727, p=0.8927. N=5 per condition. **D**. Finasteride did not affect the thermal antinociceptive effect of morphine as measured by the time before paw lick in response to different temperatures 30 minutes after treatment with morphine. Two-way ANOVA 48.5°C significant effect of Treatment F(2,9)=770)p<0.0001, 51.5°C significant effect of Treatment F(2,9)=33.12)p<0.0001. Both vehicle+morphine and finasteride+morphine are significantly different from untreated animals. No difference observed between vehicle+morphine and finasteride+morphine. N=4 per condition. \**p*-value < 0.05, ** *p*-value<0.01. error bars +/- s.e.m

After the conditioning phase, animals were separated into two groups, and the effect of finasteride was tested over 10 sessions. Each group was treated with either *an* intraperitoneal injection of finasteride (50 mg/kg) or vehicle for five out of the ten sessions. Treatments were then inversed for the remaining five sessions, so that both groups received five sessions of finasteride and five of vehicle. As observed for hydrocodone self-administration, acute treatment with finasteride significantly reduced the number of active nose-pokes (Figure 4B-C), as well as the total amount of fentanyl consumed per session (Figure S6B-E), without affecting the number of inactive nose pokes (Figure S7). This acute treatment with finasteride was effective during the five days of treatment, regardless of the order of treatment (Figure 4D-E). Importantly, finasteride reduced opioid consumption in both male and female animals (Figure S8).

We also quantified the number of active nose-pokes over time during each session. In vehicle-treated animals, there was a rapid succession of active nose-pokes in the first 10 minutes before slowing down over the next 50 minutes. Finasteride treated animals followed a similar pattern, but they only acquired approximately half of the nose-pokes before slowing their rate of intake (Figure 4F).

These results collectively demonstrate that the activity of finasteride on opioid self-administration is conserved in mammals, even for animals conditioned with the synthetic opioid fentanyl.

### Finasteride does not affect the antinociceptive properties of opioids

Despite their high abuse potential, opioids are still invaluable as analgesics, and in the absence of an effective alternative, are still an essential treatment, especially for people suffering from chronic pain. Therefore, an ideal candidate for an opioid abuse treatment would be a drug that doesn’t affect the pain-killing properties of opioids. A previous report demonstrated that finasteride did not reduce the efficiency of morphine; however, the authors used a lower dose of finasteride (25mg/kg) and tested a single nociceptive stimulus^39^. To further validate that finasteride doesn’t interfere with the antinociceptive effects of opioids in our conditions, we used a rat model of neuropathy.

We performed spinal nerve ligation (SNL) surgery on rats^40^, and fourteen days after surgery, animals were separated into different groups. Mechanical allodynia and nociception were measured utilizing von Frey Hair and Randall-Selitto, respectively^41,42^. We also measured thermal nociception (hot plate)^43^. We first validated the antinociceptive effects of opioids by testing different doses of morphine (1-3mg/kg, SC) or hydrocodone (1-10mg/kg, SC) every 30 minutes for 3 hours. For Randall-Selitto, an increase in mechanical force is applied to the paw until a withdrawal response is observed (Figure 5). Treatment with either morphine or hydrocodone significantly increased the force needed to trigger withdrawal compared with the same animals prior to treatment with the opioid (Figure 5B and Figure S9A). We then selected the most efficient dose for both opioids, morphine (3 mg/kg), and hydrocodone (10 mg/kg) and tested the effect of co-injection with finasteride (50 mg/kg, IP). The antinociceptive effect of neither opioid was reduced by the co-treatment with finasteride (Figure 5C and Figure S9B). We also performed the same set of experiments to test a different mechanical stimulus, i.e., the Von Frey test. As with the Randall-Selitto test, we did not detect any reduction in the antinociceptive effects of the opioids by treatment of the rats with finasteride (Figure S10).

In order to test a different type of nociception, we used the hot plate assay to test for a thermal stimulus on a different group of neuropathic rats. Animals were placed on a hot plate analgesia meter and the latency to lick their left hind paw was measured at different temperatures (48.5°C and 51.1°C). We selected the most effective dose of opioid based on the Randall-Selitto test and performed the assay 30 and 60 minutes after opioid injection. The latency for the first lick or paw retraction of animals treated with morphine (3 mg/kg) was significantly increased at both temperatures after 30 minutes (Figure 5D) and 60 minutes (Figure S11) when compared to untreated animals. Interestingly, treatment with hydrocodone (10 mg/kg) did not reduce the latency at 51.5°C and was not very effective when given 60 minutes prior to the test. As with mechanical nociception, co-injection with finasteride (50 mg/kg, IP) did not reduce the antinociceptive efficiency of either opioid at any temperatures or pretreatment times (Figure 5C and Figure S11). The fact that finasteride does not inhibit the antinociceptive or antiallodynic effect of either morphine or hydrocodone in response to different painful stimuli in a rat model of neuropathic pain demonstrates that finasteride is unlikely to interfere with the principal beneficial activity of opioids and might, therefore, be used to reduce abuse liability while retaining the clinical utility of opioids.

### Steroids regulate opioid self-administration

Because finasteride is a known 5αR inhibitor, we hypothesized that finasteride reduces opioid self-administration by altering the level of one or more neuroactive steroids in the brain. To identify candidate neuroactive steroids mediating this effect, we isolated whole brains from wild-type zebrafish and from opioid-conditioned zebrafish treated with either DMSO or finasteride (10 µM). We then performed steroid quantification using untargeted ultra-performance liquid chromatography-mass spectrometry (UPLC-MS).

We normalized the results (min-max) for each steroid and compared the levels between DMSO- and finasteride-treated conditioned animals. The only steroid that reached significance by itself was dehydroepiandrosterone sulfate (DHEAS), which was markedly increased in finasteride-treated animals (Figure 6A). We also observed that other precursor steroids, including testosterone and pregnenolone, also showed the same trend of accumulating in finasteride-treated animals. Surprisingly, we also detected a reduction in dehydroepiandrosterone (DHEA), which could suggest an increase in the conversion of DHEA to DHEAS in treated animals. Interestingly, we did not detect the same trend for other sulfated steroid species (Figure S12). The opposite trend of decreases in finasteride-treated animals was also observed for steroids downstream of 5αR, especially allopregnanolone (AP) and 3α-androstanediol, but these differences did not reach statistical significance (Figure 6A-B).

**Figure 6.**
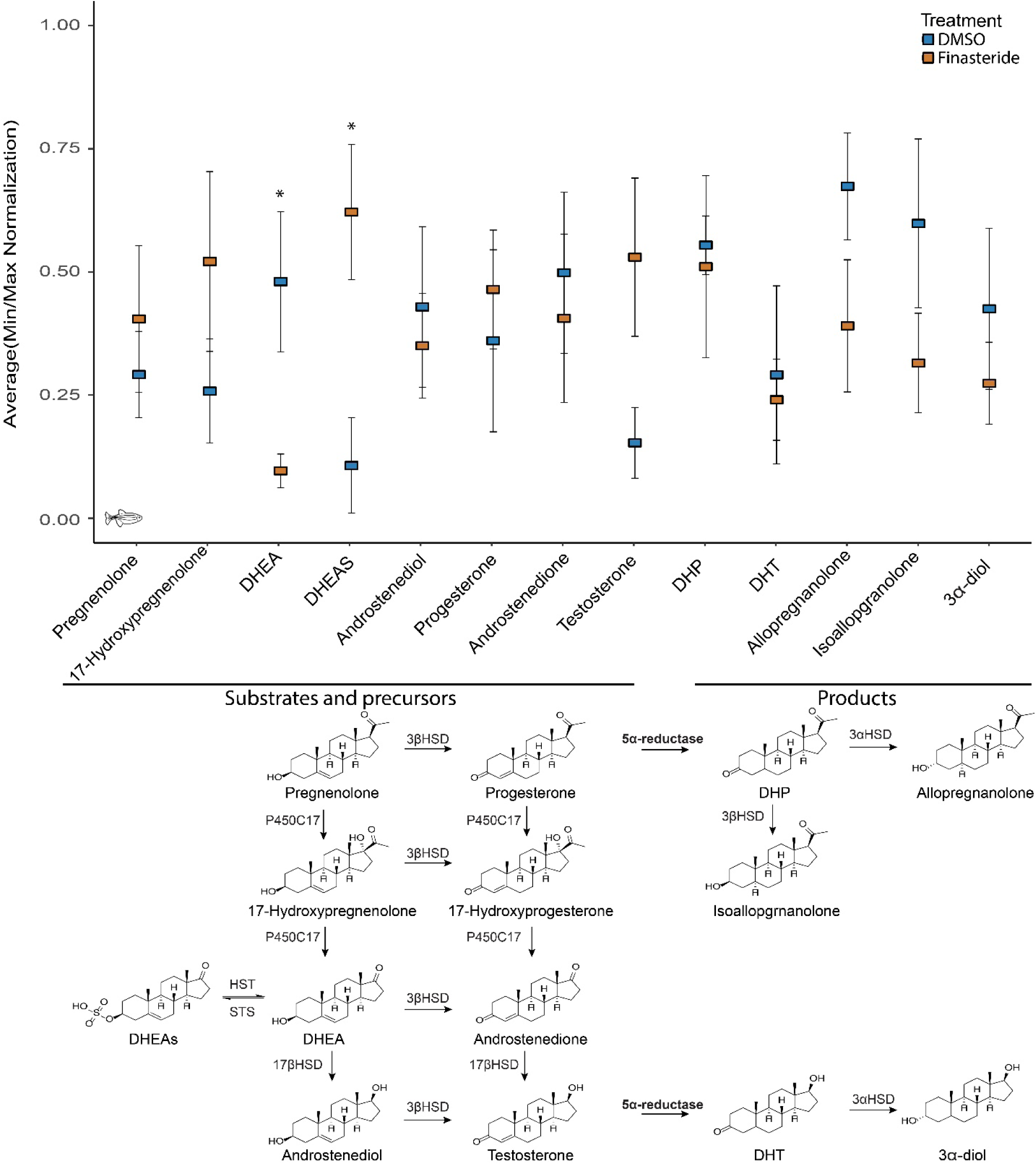
Finasteride treatment changes neurosteroid levels in the conditioned animal brain. **A**. Normalization score for the quantification of steroids in conditioned brains treated with DMSO or finasteride (10 µM). *p*-value calculated with Student’s T-test. n=5 set of 10 brains per condition. \**p*-value<0.05 **B**. Partial neurosteroidogenesis pathways. Finasteride blocks the rate limiting enzyme, 5α-reductase, causing accumulation of upstream neurosteroid species.

We tested how the accumulation of DHEAS affects opioid self-administration. Given that sulfated steroids are less effective at crossing the blood-brain barrier^44,45^, we incubated conditioned zebrafish with 10 µM DHEA for 60 minutes before measuring opioid-self administration. As observed with finasteride, incubation with DHEA drastically reduced the number of visits at the active platform (Figure 7A). We also tested the primary DHEA precursor, pregnenolone, and again observed a reduction in opioid self-administration (Figure 7A). Taken together, these results suggest that the accumulation of DHEAS and other precursors observed in finasteride-treated animals could play an important role in the reduction of opioid self-administration. The fact that DHEA-treated animals had a reduced number of visits at the active platform also suggests that the reduction in DHEA level observed in finasteride-treated animals is a result of an increased conversion to DHEAS.

**Figure 7.**
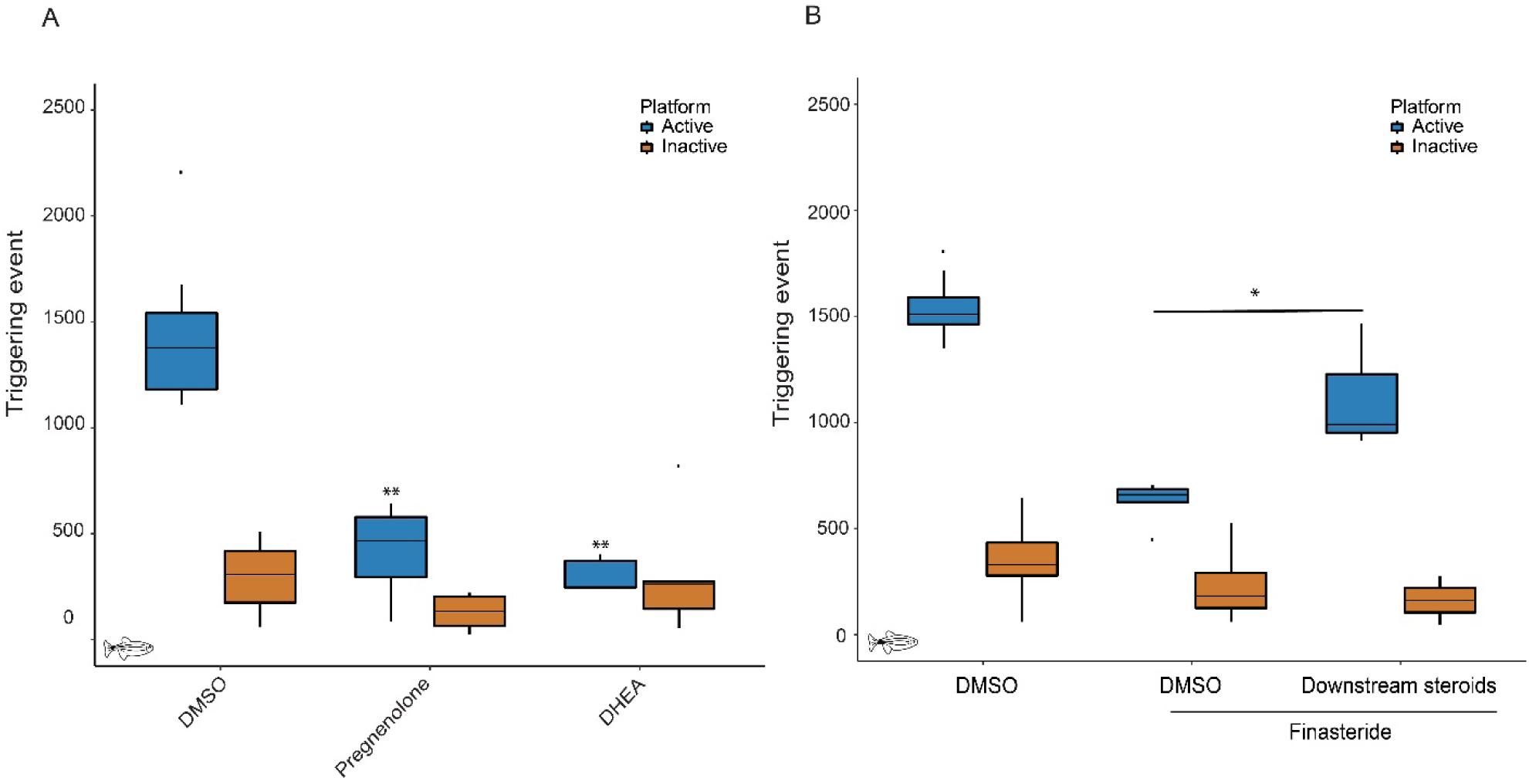
Specific neurosteroids also affect opioid self-administration. **A**. Incubation with steroids upstream of 5αR, DHEA (10µM) and pregnenolone (10µM) reduces the number of triggering events at the active platform. *p*-value computed by Tukey HSD on one-way ANOVA, Inactive platform (F(2,21)=2.175)p=0.139 and Active platform (F(2,21)=51.09)p=8.57E-09. No significant difference was detected for the inactive platform compared to the DMSO control. DMSO n=12, Pregnenolone n=8, and DHEA n=5. **B**. Co-treatment with finasteride (10 µM) and a selection of steroids downstream of 5αR partially blocks the reduction in opioid self-administration induced by finasteride. *p*-value computed by Tukey HSD on one-way ANOVA, Inactive platform (F(2,16)=1.575)p=0.239 and Active platform (F(2,16)=68.93)p=1.37E-08. No significant difference was detected for the inactive platform. DMSO n=8, finasteride n=8, finasteride+downstream steroids n=3. \**p*-value < 0.05, ** *p*-value<0.01. Each n represents a group of 15 animals.

Since we detected a reduction in some products of 5αR after treatment with finasteride, we decided to test whether their reintroduction could interfere with the activity of FIN. However, since we did not detect any significant changes for a single product, we decided to test if a combination of steroids from the same class could act together. We chose to co-treat conditioned animals with finasteride (10µM) and allopregnanolone (0.1µM), androsterone (1µM) and 3α-diol (1µM) for 60 minutes. Interestingly, the presence of these products was sufficient to partially block the effect of finasteride and significantly increase the number of visits at the active platform when compared to finasteride treatment alone (Figure 7B).

Taken together, these results suggest that an accumulation of DHEA or its sulfated form might play a critical role in mediating the effect of finasteride. However, other neurosteroids such as 5αR products may also be involved in the regulation of opioid self-administration.

## Discussion

We previously described the development of an assay to condition zebrafish to self-administer the opioid hydrocodone, representing a new tool for OUD research. One of the main advantages of using zebrafish is the possibility of systematically screening drugs for therapeutic effect in a whole organism^46^. We thus performed a small-scale small molecule screen to identify drugs that reduce opioid self-administration. One of the most effective compounds was the 5α-reductase (5αR) inhibitor finasteride, which substantially reduces opioid self-administration but does not affect food self-administration in fish, suggesting that it does not affect motivated behaviors generally. This observation also confirms that treated animals can still recall and find the location of the active platform in the arena. Consistent with this idea, it is also noteworthy that we did not detect significant differences in the number of triggering events at the inactive platform between the control and the finasteride-treated fish.

Although the zebrafish self-administration model enables screening of candidates at higher scale, rodent models are more established in the field, so we sought to confirm the effect of finasteride in mammals by using rat models. As with zebrafish, acute treatment with finasteride is sufficient to reduce self-administration of two different opioids conditioned with two different paradigms in rats, indicating that the effects of finasteride on opioid self-administration are robust and broadly conserved.

Although finasteride is typically used for treating non-neuronal indications, there is evidence to suggest it might also have beneficial effects in the nervous system. In animal models, inhibition of 5αR by finasteride has been shown to improve symptoms caused by pathologies driven by dopamine imbalances^47,48^. Dopamine is a central component of the reward pathways, and increased dopaminergic activity has been linked to drug abuse^49^. Also, a recent study revealed that rodents treated with finasteride had reduced reactivity towards both incentive stimuli and stress responses. There is a growing body of evidence that an increase in neuronal stress pathways plays an important role in substance abuse disorders^50,51^. Taken together, these studies indicate finasteride may have an impact on key neuronal pathways linked to OUD.

Finasteride is an inhibitor of key enzymes in steroid production, 5α-reductases. The three 5α-reductase isoenzymes SRD5A1, SRD5A2, and SRD5A3 are expressed in different tissues, including the nervous system^52–56^. In zebrafish, we detected multiple isoenzymes in the zebrafish brain, and the transcripts are present in different cell types. In other animal models, the inhibition of 5αR leads to major changes in steroid levels where the levels of steroid species upstream of 5αR such as progesterone or testosterone increase. In contrast, the levels of products of 5αR and downstream steroids are decreased^21,22,57^. Because the effects of finasteride on the steroid profile are pleiotropic, characterization of the specific steroid species involved in the regulation of opioid self-administration may eventually lead to the development of even more precise, targeted therapies for OUDs.

To investigate the landscape of changes induced by the treatment of opioid conditioned animals with finasteride, we used an untargeted approach to quantify steroids in the brain. In other models, finasteride induces changes in steroid levels in specific brain regions^58,59^, but because of the small size of the zebrafish brain, we used whole brains and could not achieve the same level of regional specificity. Nonetheless, we identified interesting trends, including an accumulation of several 5αR precursors and a reduction in 5αR products after treatment with finasteride.

Our results suggest that accumulation of DHEAS plays an important role in opioid self-administration. It is unclear why we detected a decrease in the non-sulfated DHEA levels and an accumulation of DHEAS after treatment with finasteride. This effect appears to be specific for these two steroids as we did not detect significant accumulation of other sulfated steroids. There are different steroid sulfotransferases involved in steroid sulfation in humans, where DHEAS is formed by SULT2A1 and SULT1E1, but pregnenolone sulfate is formed by SULT2A1 and SULT2B1a ^60,61^. It has been shown that zebrafish express putative sulfotransferases with similarities to human SULT2B1b and SULT2A1. However, the conservation of SULT1E1 has not been demonstrated^62,63^. As with humans, the enzyme involved in the formation of sulfated steroids may be different depending on the substrate. This could explain why only DHEAS is significantly accumulating in zebrafish brains upon finasteride treatment. Other precursors, such as pregnenolone and testosterone, also show some degree of accumulation; thus, the accumulation of DHEAS may be caused by a compensation mechanism driven by a higher rate of conversion of DHEA to DHEAS. Another possibility that could explain the accumulation of DHEAS is an effect on steroid sulfatase (STS) activity, which is responsible for the conversion of DHEAS back to DHEA. However, as opposed to sulfotransferases, there is a single enzyme performing this function for all steroids^63^. Therefore, the absence of effects on other sulfated steroids suggests that the accumulation of DHEAS is likely not a result of an inhibition of STS.

The fact that incubation with DHEA can reduce opioid intake further supports the hypothesis that DHEA/DHEAS plays an important role in opioid intake regulation. These neuroactive steroids have been shown to affect neuronal activity by binding to different neurotransmitters receptors including key pathways relevant to substance abuse such as glutamatergic (NMDA), GABAergic and dopaminergic systems^64–66^. The modulation of these pathways has been shown to be important in the regulation of opioid abuse disorders^18,49,67–71^ and could explain why incubation with DHEA reduces opioid intake. Furthermore, previous reports describe a link between DHEAS and the development of opioid tolerance without showing an effect on self-administration^72^. Additionally, chronic treatment with DHEA has also been shown to reduce cocaine self-administration and reinstatement in rats^73^.

Although we believe that DHEA/DHEAS plays a key role in the effect of finasteride on opioid self-administration, we also have evidence that other steroid species could be involved. Similarly to DHEA, these other steroids have been shown to affect key neuronal pathways relevant to substance abuse involving GABAergic and dopaminergic systems ^74–78^. Some of these steroids, such as allopregnanolone, have been linked to neuronal stress response^79,80^. Therefore, by reducing allopregnanolone production, finasteride may reduce the negative affective state that contributes to opioid self-administration.

We currently do not know if finasteride influences opioid self-administration by reducing motivation for drug seeking or if treated animals are simply satisfied with a lower amount of drug. Our results in the fentanyl self-administration assay revealed that finasteride treated rats initially perform nose-pokes to self-administer fentanyl at a rate similar to their untreated controls, but they slow down their intake sooner than controls, suggesting finasteride-treated animals may achieve satiation with a lower amount of fentanyl. A more detailed study on the impact of finasteride on OUD will be required to further validate that finding.

We have demonstrated that finasteride modulates opioid consumption, a critical aspect in OUD, in two different animal models, and for different classes of opioids. If these results are translated in humans, repurposing finasteride for the treatment of opioid abuse may be a viable therapeutic strategy to treat OUD. Finasteride was previously found to reduce risk-taking behavior, a behavioral feature typically associated with substance use disorders^48,81,82^. Furthermore, preliminary observations suggest that finasteride may help reduce pathological gambling^83^. This finding further supports finasteride’s potential in the treatment of addictive behavior. Since finasteride is FDA-approved and its side effects and toxicology are well-studied, these results could rapidly lead to clinical trials to test the therapeutic potential of this drug for OUD. These results suggest that finasteride may help patients control their drug intake, which could mitigate the risk of overdose.

Finasteride has been clinically used since the 1990s for the treatment of androgenic alopecia and benign prostate hyperplasia. Although it is considered a well-tolerated and relatively safe drug, there is evidence of sexual disfunction in 3.4 to 15.8 percent of men. A rare but serious side effect known as post-finasteride syndrome (PFS) has also been reported^84–86^. PFS prevalence is unclear but it manifests as a range of persistent physical and neuropsychiatric disorders such as depression and anxiety that develop during or after discontinuation of finasteride use. Clinical studies will be needed to understand the side-effects associated with the use of finasteride for the treatment of OUDs, and careful clinical consideration must be given in weighing potential risks and benefits of finasteride use.

The fentanyl self-administration study presented here reveals that both male and female rats respond to treatment with finasteride. These data raise the possibility that finasteride could be used as a treatment for OUD in both males and females. This finding is interesting, given that finasteride is mainly used in men (although there is some history of use to treat hirsutism in women)^87^.

An ideal treatment for opioid abuse would not block opioid antinociceptive action. It has been shown that finasteride does not block the antinociceptive effect of morphine^39^. To extend these findings, we used the most rigorous test available in rats by testing the effect of opioids on a rat model of neuropathy. We performed two different types of tests to measure nociception: mechanical and thermal. Testing multiple different assays in a sensitized background confirmed that finasteride does not affect the antinociceptive effects of either morphine or hydrocodone. Our findings suggest that finasteride could potentially be used to prevent opioid abuse while still allowing opioids to be used when necessary for pain relief.

In conclusion, the present study identifies the widely-used drug finasteride as an effective agent for reducing opioid intake across multiple species and self-administration paradigms. The data further indicate that the DHEA/DHEAS pathway is a major mediator of finasteride’s effect. These findings point to a promising potential therapeutic strategy for treating OUD and open new avenues for investigating the role of specific steroids in regulating opioid use behaviors.

## Materials and Methods

### Zebrafish

#### Fish handling

Adults zebrafish (*Danio rerio*) of the wild type Ekkwill strain were maintained in the fish facility at 28–29°C with a 14/10 hours light/dark cycle. All zebrafish experiments were approved by the University of Utah Institutional Animal Care and Use Committee.

#### Adult fish treatment

Adult fish were transferred to a small treatment chamber (USplastic, USA) with 100 mL of fish water and the compound of interest was injected directly into the water. Fish were allowed to swim in the treatment solution for one hour prior to the self-administration assay.

#### Zebrafish self-administration

The same protocol as detailed in Bosse et Peterson, 2017*(89)* was used to condition fish in small groups of 15 animals. For the screen, fish were conditioned for 4 days and treated with the different compounds on day 5, before being tested in the arena for 30 minutes. For opioid conditioning, we used the following doses: hydrocodone and morphine 6 mg/L and fentanyl 0.12 mg/L. Opioids were diluted in fish water. A large number of animals were conditioned simultaneously and randomly assigned to different treatment conditions.

#### Chemicals used

Finasteride (Sigma-Aldrich, USA), Hydrocodone bitartrate (Spectrum Chemicals, USA and NIH), Morphine (Spectrum chemicals), Fentanyl (Spectrum Chemicals, USA), Dutasteride (Cayman chemical company, USA), Pregnenolone (Sigma-Aldrich, USA), DHEA(Sigma-Aldrich, USA), Methadone (Sigma-Aldrich, USA), Allopregnanolone (Tocris, USA), 3α-diol (Steraloids, USA), Androsterone (Steraloids, USA), Fentanyl (Sigma-Aldrich, USA). Compounds were either resuspended in DMSO or water according to manufacturer recommendation.

#### Food conditioning

The same apparatus was used as for hydrocodone conditioning. For food conditioning, fish were trained directly in the arena without performing the pre-conditioning protocol. Fish were trained for 50 minutes daily in a small group. Larvae food Ziegler #4 (VWR, USA) was suspended in fish water.

#### Fish locomotion

A high-speed infrared camera (Point Grey, Canada) was used to record 1-minute videos of animals in the self-administration arena. Zebrafish tracking and movement quantification were performed with the software Actualtrack (United Kingdom).

#### Statistical analysis

For zebrafish self-administration data, R graphic programming was used to generate the plots. ANOVA tests were run on plot data to test significance. ANOVA tests were performed first on inactive platforms for each dataset to validate that there was no difference between the different conditions and then active platform values were used to test for significance. All boxplots were generated using R graphic programming and the *ggplot* module. The lower and upper hinges correspond to the first and third quartiles. The line is the median. The whiskers extend from the hinged to the maximum or minimum value at most 1.5x the inter-quartile range (IQR) from the hinge. Data points beyond that are considered outliers. No data points were excluded from the statistical analysis.

#### Single cell data analysis

Processed single cell data from *Raj et al. Nat Biotechnol(90)* from 6 individual whole zebrafish brains (f1 n=6,759, f2 n=7,112, f3 n=15,156, f4 n=12,121, f5 n=9,919, f6 n=6,009) as well as manually dissected fore- (n=3,615), mid- (n=1,504) and hindbrains (n=3,894) were downloaded as an R data object from Gene Expression Omnibus (GSE105010). The R package Seurat (v3.1.1) was used to create a new Seurat object from the raw counts data found within the downloaded InDrops object. Samples with less than 200 genes detected were already removed from these data, but we additionally removed outliers with more than 6000 genes detected and those higher than 25% mitochondrial genes. This further processing removed an additional 371 samples. We followed the SCTransform workflow as recommended in the Seurat vignette (https://satijalab.org/seurat/v3.1/sctransform_vignette.html) for scaling, normalization and finding variable genes. Mitochondrial mapping percentage was regressed during normalization. A principal component analysis was first performed, then clustering using the Seurat FindNeighbors (using the first 30 dimensions) and FindClusters function (resolution of 2.5). The data were visualized using t-SNE dimensionality reduction using the first 30 dimensions. Similar to the published analysis in *Raj et al. Nat. Biotechnol*., we observed that the 65,718 samples produced 61 clusters. t-SNE coordinates, expression values from the SCT “data” slot and metadata were exported from the Seurat data object to create t-SNE figures using data.table and ggplot2 demonstrating the cluster identities, broad brain regions (fore-, mid- and hindbrain) and srd5a family expression. We plotted all family members highlighting only those cells in the foreground with relatively medium to high expression values where expression cut-offs were determined for each marker individually based on expression distribution (srd5a1 0.8-3.0; srd5a2a 1.0-3.0; srd5a2b 1.0-3.0; srd5a3 0.8-3.0)^34,88–93^.

#### Steroid quantification Brain extraction

Conditioned fish were transferred to a small treatment chamber (USplastic, USA) with 100 mL of fish water and treated with either DMSO (0.02%) or finasteride (10µM). Fish were allowed to swim in the treatment solution for one hour. Treated animals were then transferred to a water bath with ice-water for euthanasia. The brain of each animal was then extracted in PBS 1X. The head was cut using a razor blade behind the gill, the skull was then carefully peeled to expose the brain using surgical forceps. The brain was then extracted by performing a cut at the base of the cerebellum. The extracted tissue was then placed in 1.5 mL self-standing microcentrifuge tube, (USAscientific, USA) on ice and the brains of 10 animals were pooled together in the same tube. Any liquid was then removed from each tube before weighing the tissues. Samples were then flash frozen in liquid nitrogen and placed at −80°C until extraction.

#### Chemical

Reference standards were purchased from Steraloids (Newport, USA). All solvents were HPLC grade, and all other chemicals used were of the highest grade available. Stock neurosteroid standard mixture was prepared by mixing 5 µL of 1mg/mL solution of each steroid and adjusting the final volume to 1mL by using methanol*(97-99)*. All the stock solutions were stored at −80°C.

#### Sample preparation

Tissue samples were extracted as described previously*(97-99)*. Briefly, tissue samples were extracted with 1 mL chloroform. The mixture was vortexed for 30 sec and centrifuged for 5 min; the chloroform layer was transferred to 2 mL tube and dried. The resulting residue was extracted with 1 mL MeOH. The MeOH layer mixture after 5 min centrifugation was added to the above chloroform extract. This mixture was dried and re-suspended in 125 µL MeOH and filtered using 5kD membrane filters. Filtrates were transferred to vials for UPLC-MS analysis.

#### UPLC-MS Analysis

Tissue sample extracts were subjected to UPLC-MS analysis for the measurement of neurosteroids, as described previously*(97-99)*. UPLC analyses were carried out using a Waters Acquity UPLC system connected with the high-performance triple quadrupole mass spectrometer. Analytical separations on the UPLC system were conducted using an Acquity UPLC C18 1.6 µ column (2.1 x 150 mm) at a flow rate of 0.15 mL/min and C18 1.7 µ column (2.1 x 50 mm) at flow rate 0.2mL/min. For the first column, the gradient was started with 100% A (0.1% formic acid in H2O) and 0% B (0.1% formic acid in CH3CN), after 0.1min changed to 80% A over 1 min, and then 45% A over 5min, followed by 20% A in 2min. Finally, over 0.5 min, it was changed to 0% A, then after 13 min, it was changed to the original 100% A over 1 min, resulting in a total separation time of 13 min. For the second column, the gradient was started with 100% A (0.1% formic acid in H2O) and 0% B (0.1% formic acid in CH3OH), after 0.1min changed to 80% A over 2 min, and then 45% A over 2 min, followed by 20% A in 2min. Finally, over 1 min, it was changed to 0% A, then after 8 min, it was changed to the original 100% A over 2 min, resulting in a total separation time of 10 min. The elution from the UPLC column was introduced to the mass spectrometer. All MS experiments were performed by using electrospray ionization (ESI) in both positive ion (PI) and negative ion (NI) mode, with an ESI-MS capillary voltage of 3.5 kV, an extractor cone voltage of 3 V, and a detector voltage of 650 V. The following MS conditions were used: desolvation gas at 400 l/h, desolvation temperature at 350°C and source temperature 150°C. Pure standards of all targeted neurosteroids were used to optimize the UPLC-MS/MS conditions prior to analysis and performing calibration curves ^94–96^. Reference standards were run before the first sample, in the middle of the runs and after the last sample to prevent errors due to matrix effect and day-to-day instrument variations. In addition, spiked samples were also run before the first sample and after the last sample to calibrate for the drift in the retention time of all neurosteroids due to the matrix effect. After standard and spiked sample runs several blanks were injected to wash the injector and avoid carry-over effects. Resulting data were processed by using Target Lynx 4.1 software (Waters) ^94–96^.

#### Data normalization

Steroids counts were first normalized using the initial weight of the tissue before extraction. To compare the levels of the steroids, we then used min-max normalization for steroid count in each sample.

### Rats

#### Hydrocodone self-administration and nociception Animals

Adult male Sprague-Dawley rats (Charles River, Roanoke, USA) were group-housed (3-4/cage) within rooms maintained at 22 ± 2 °C and 55% humidity, on a 12/12 h light/dark cycle (lights on at 7:00 AM). Food and water were available *ad libitum*. Following a 7-day acclimation to the housing facilities, animals were handled daily for 5 min. Behavioral measurements were carried out and analyzed by trained experimenters in a blinded fashion.

#### Chemicals

Hydrocodone (Spectrum Chemical, USA) and morphine (Spectrum Chemical, USA) were dissolved in a solution of 2% DMSO and 98% saline. Finasteride (Astatech, Bristol, USA) was suspended in a solution of 5% DMSO, 5 % Tween 80 and 90% saline (5:5:90).

#### Hydrocodone self-administration

Apparatus: The apparatus consisted of 8 operant conditioning chambers (Habitest, Coulbourn, USA), measuring 30.48 cm (W) x 25.4 cm (D) x 30.48 cm (H), and enclosed in sound-attenuating cubicles with ventilation fans. Each chamber was equipped with two retractable levers: an active lever coupled to the intravenous delivery of hydrocodone, and a control (inactive) lever. Active lever placement on the left or right side followed a counterbalanced order. Three cue lights were placed over the active lever. The apparatus was controlled by Graphic State 4 software (Coulbourn, USA).

Experimental procedure: Opioid self-administration was performed using a modified version of the protocol described by *Mavrikaki et al*^97^. Rats weighing 225-250 g were used. Rats were anesthetized with ketamine and xylazine and underwent catheterization surgery. Briefly, a polyurethane catheter was inserted through the external jugular vein, passed under the skin, and fixed in the mid scapular region. Post-operative care included buprenorphine and enrofloxacin for analgesic and antibiotic management, respectively. Catheter patency was maintained through daily flushing with a heparin (500 IU/mL) / 50% dextrose solution.

Ten days after surgery, all rats were gently handled and kept under a food restriction regimen that maintained them at 90% of their initial body weight and was continued throughout the whole behavioral procedure. A syringe containing a hydrocodone solution was placed in an infusion pump located outside the chamber and connected to the rat’s catheter via a fluid swivel and spring-covered Tygon tube suspended through a counterbalanced swivel. The solution was administered at a dose of 0.016-0.128 mg/kg/infusion, in a volume of 160 μL/kg/infusion. Operant training began three days later and consisted of three stages of fixed-ratio reinforcement schedule: FR1, FR2, and FR5. Rats underwent daily, 1 h-long experimental sessions, between 9:00 AM and 3:00 PM and for 7 days/week, consisting of a sequence of trials (Figure 3). Each trial began with a 5-s period, during which the house light was turned off and the cue light blinked three consecutive times. Subsequently, the house light was turned on and both levers were extended. Once the rat completed the fixed ratio on either lever, both levers retracted, and a new trial began after a 15-s time-out period. Animals progressed to the next stage after reaching > 70% of total lever presses being on the active lever. Animals reached FR5 stability after 31 days of training and were then treated with either finasteride or its vehicle.

#### Locomotor activity

Rats were tested for locomotor activity in an open field surrounded by black Plexiglas walls (47 cm × 47 cm × 47 cm). Comparisons were drawn between self-administering animals (immediately after their performance in the operant chamber) and control animals not receiving any opioids. The locomotor analysis was performed using EthoVision XT 14 pathway tracking software (Noldus Instruments, Wageningen, The Netherlands).

#### Neuropathic pain assessment

Experimental procedure: Sprague-Dawley rats weighing 150-180 g were used. Following 2-3 days of handling, rats underwent mechanical nociception testing via von Frey Hair and Randall-Selitto tests. Neuropathy was then induced by SNL surgery, as previously described^40^. Rats were anesthetized using xylazine and ketamine (10/75 mg/kg, IP), and their left L5 spinal nerve was exposed and tightly ligated with 4.0 silk suture (Mersilk^®^, Ethicon thread, Johnson). Muscle, fascia, and skin were then sutured, and the rats were treated with enrofloxacin (10 mg/kg, SC) and carprofen (5 mg/kg, SC) for post-operative care. Fourteen days after surgery, nociception was re-tested, and allodynia was confirmed in rats exhibiting a >30% reduction of their pain threshold. Rats were then assigned to different treatment groups to receive either morphine (1-3 mg/kg, SC), hydrocodone (1-10 mg/kg, SC) or saline. The antinociceptive effects of opioids were tested every 30 min for 6 consecutive observations. The analgesic effects of morphine and hydrocodone (at their most effective doses) were also tested in combination with finasteride (50 mg/kg, IP) or its vehicle, to ascertain whether finasteride altered the antiallodynic properties of opioids. The effects of opioids and finasteride were also tested for thermal nociception in a separate group of rats with spinal nerve ligation using the hot plate procedure.

von Frey Hair Test: Tactile allodynia was assessed using a set of 8 von Frey monofilaments (Bioseb, Vitrolles, France) with logarithmic incremental stiffness (of 1.4, 2, 4, 6, 8, 10, 15 and 26 g). Paw-withdrawal threshold was measured, and 50% response threshold was calculated using the Up-Down method and Dixon’s formulae, as previously described^41^. Behavioral assessments were run prior to and 14 days after SNL surgery. Rats were individually placed in plexiglass compartments (17 × 11 × 13 cm) with a wire mesh bottom that allowed full access to paws. After 20-30 min of acclimation, a first 6-g hair was perpendicularly applied against the plantar surface of the left hind paw for 6 s. Paw withdrawal and/or licking reflex was considered as a positive response. Depending on the positive or negative response, the next filament with either lower or higher force was tested, respectively. Testing continued until either four consecutive negative or five consecutive positive responses were recorded after the first change of direction.

Randall-Selitto Test: Nociceptive withdrawal threshold was assessed using the Randall-Selitto algesimeter (Ugo Basile, Varese, Italy), as previously described^42^. Following daily handling and acclimation to the apparatus, rats were wrapped into a cotton cloth and immobilized, and the medial portion of the plantar surface of the left hind paw was carefully placed on the tip of the device. An increasing mechanical force was applied until a withdrawal response was observed. Paw withdrawal threshold for Randall-Selitto experiment is set at 25g of for applied. Rats were tested every 30 minutes for three consecutive hours following treatment (6 applications in total). Hot Plate Test. Thermal nociception was assessed using the hot plate analgesia meter (IITC Life Science, Woodland Hills, USA). The rat was placed on a plate maintained at different temperatures (48.5 and 51.5 °C), and their progressive latencies to lick the left hind paw were measured.

#### Fentanyl self-administration Subjects

A total of 20 adult male and adult female Wistar rats (Charles River, USA) weighing 200 – 475g at the start of the experiment were individually housed and kept on a 12-h light/ 12-h dark cycle in a temperature and humidity-controlled room. Animals were provided food and water *ad libitum*.

#### Drugs

Fentanyl Citrate (Medisca, USA) was dissolved in deionized water at a concentration of 50μg/mL. Finasteride (Astatech, USA) was suspended (50mg/mL) in a solution of 2.5% ethanol, 5% Tween80, and 92.5% saline.

#### Fentanyl self-administration

Apparatus: Oral fentanyl self-administration tasks were completed in eight modular operant chambers (Med Associates, USA), equipped with a liquid magazine tray stationed between two nose-poke devices. Additionally, the operant chamber was outfitted with a solenoid controlled liquid valve (Lee Valves, USA) and a set of audiovisual cue equipment [house light, magazine light, and a tone generator].

Experimental procedure: A operant oral self-administration behavioral assay described by Shaham and colleagues*(104)* was utilized. Rats were trained to obtain liquid fentanyl delivered into a liquid magazine tray following an operant response on an FR1 reinforcement schedule. During the self-administration sessions, a nose-poke in the active port (counterbalanced between animals) resulted in fentanyl delivery (0.02mg/kg/delivery). Concurrent with drug delivery was a 10s audiovisual conditioned stimulus (CS) comprised of a 1s illumination of a light inside the nose-poke port, a 10s tone, and a 10s illumination of a light stationed above the liquid magazine tray. Any additional nose-pokes during the 10s CS were without consequence. Drug availability at the start of each session and following CS presentations was signaled by illumination of the house-light placed on the wall opposite of the nose-poke ports. All nose-pokes in the inactive port were without consequence. Animals were given two 30-minute magazine training sessions, during which any active responses resulted in fentanyl and CS presentation; however, if the animal made no responses within 2-3 minutes of the last drug delivery (or start of the session) a non-contingent fentanyl and CS delivery occurred. Following these two training sessions, animals had 15, 1-hour sessions to self-administer fentanyl for 5 days/week. During these 15 sessions, all animals reached a response criterion of >70% of nose-pokes occurring at the active nose-poke port. To test the effect of finasteride on fentanyl consumption animals received 10 additional self-administration sessions across 12 days, in which finasteride (50mg/kg, 1mL/kg) or vehicle (1mL/kg) was administered intraperitoneally 45 minutes prior to the session. Each treatment (vehicle or finasteride) was given for 5 consecutive days, with a 48-hour period before they received the opposite treatment. The order of treatment administration was counterbalanced across animals. Following self-administration sessions, mg/kg of fentanyl consumed was calculated for each animal by subtracting the amount of fentanyl left in the magazine tray from the total amount of drug delivered.

#### Study approval

All animal studies were approved by local Institutional Animal Care and Use Committees (IACUC). All zebrafish experiments were approved by the University of Utah Institutional Animal Care and Use Committee. Hydrocodone self-administration studies and nociception studies in rats were compliant with the National Institute of Health guidelines and approved by the IACUC of the University of Utah. The fentanyl self-administration studies in rats were conducted under the guidance and permission of the Institutional Animal Care and Use Committee at the University of Washington and pursuant to federal regulations regarding work with animals.

## Author contribution

G.D.B designed the experiments and performed the zebrafish assays, analyzed the data and wrote the manuscript with R.T.P. and M.B. R.C. designed the nociceptive experiments in rats and performed the experiments. G.F. designed and performed the hydrocodone self-administration assay in rats assisted by E.V. T.Z. contributed to experiment design. R.T.P. designed and supervised zebrafish experiments. M.B. designed, analyzed and supervised the experiments on rat hydrocodone self-administration experiments and nociception. N.W.G. performed steroid extraction and quantification R.D.F. J.S.L designed and performed the Fentanyl self-administration experiment in rats. R.D.F and P.E.M.P. designed and analyzed the Fentanyl self-administration in rats. All authors contributed to data interpretation and commented on the manuscript.

## Acknowledgments

We thank the NIDA Drug Supply Program for providing hydrocodone bitartrate powder. We thank C. Herdman for helpful comments and bioinformatic support. We would like to acknowledge the Centralized Zebrafish Animal Resource (CZAR) at the University of Utah for providing Zebrafish husbandry, laboratory space, and equipment to carry out portions of this research. Expansion of the CZAR is supported in part by NIH grant # 1G20OD018369-01. This work was supported by the Charles and Ann Sanders Research Scholar Award, the L.S. Skaggs Presidential Endowed Chair, and NIH grant R21DA049530 (to M.B. and R.T.P). Oral rodent fentanyl supported by National Institute of Drug Abuse (R01-DA039687. G.D.B. was supported by the Canadian Institutes of Health Research (CIHR) fellowship program. R.D.F. was supported by NIDA (T32-DA007278). T.Z. was supported by NHGRI (T32 HG008962).

