## Supplemental Figures for "The 5α-reductase inhibitor finasteride reduces opioid self-administration"

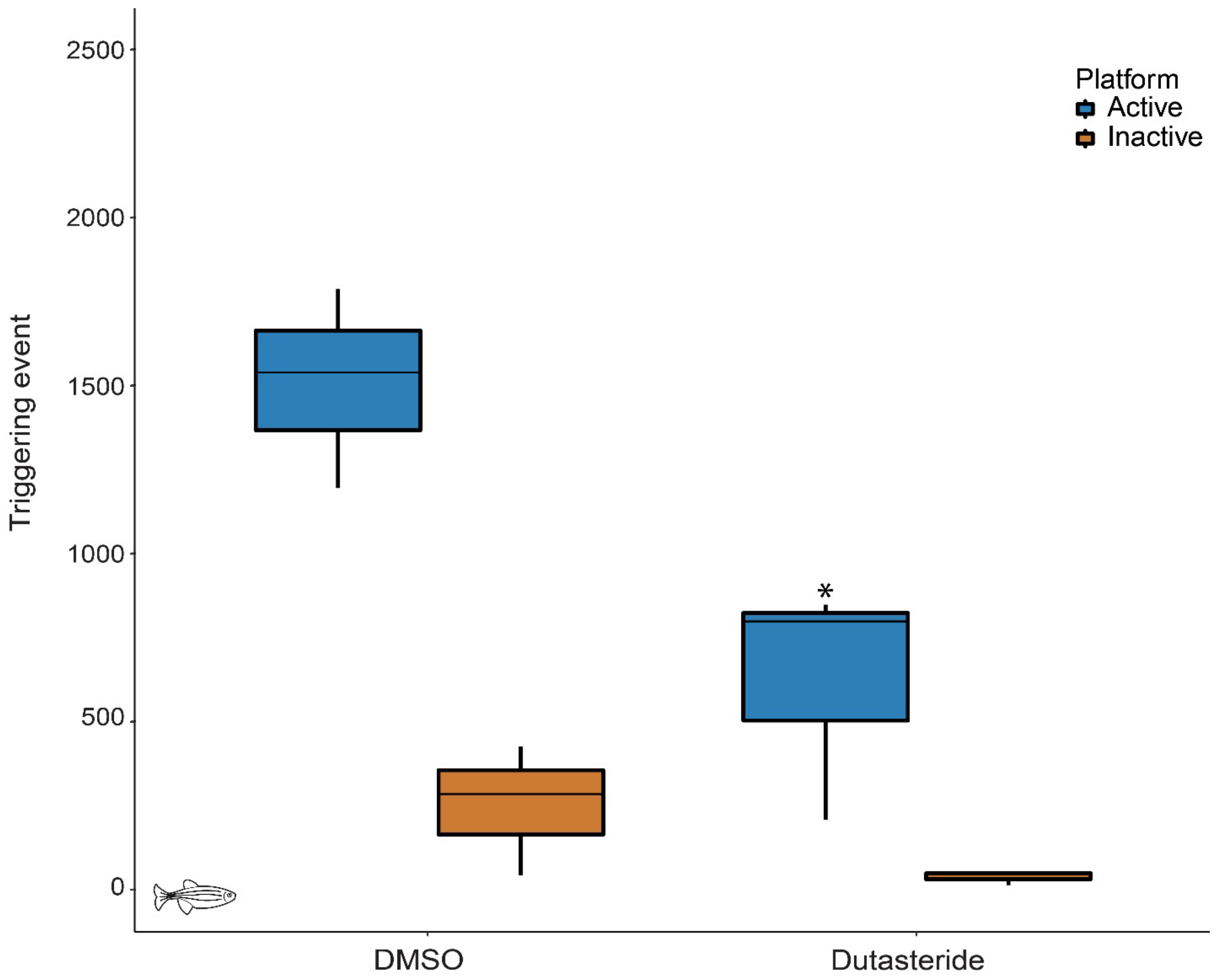


**Figure S1: Dutasteride reduces opioid-self-administration**

An additional 5αreductase inhibitor, dutasteride(10µM) also reduces the number of triggering events at the active platform. *p*-value computed by TukeyHSD on one-way ANOVA , no significant difference was observed at the inactive platform($F(1,4)=3.65$)$p=0.13$ and reach significance for the active platform($F(1,4)$,11.01)$p=0.03$. DMSO n=3, dutasteride n=3. **p*-value < 0.05. Each n represents a group of 15 animals.

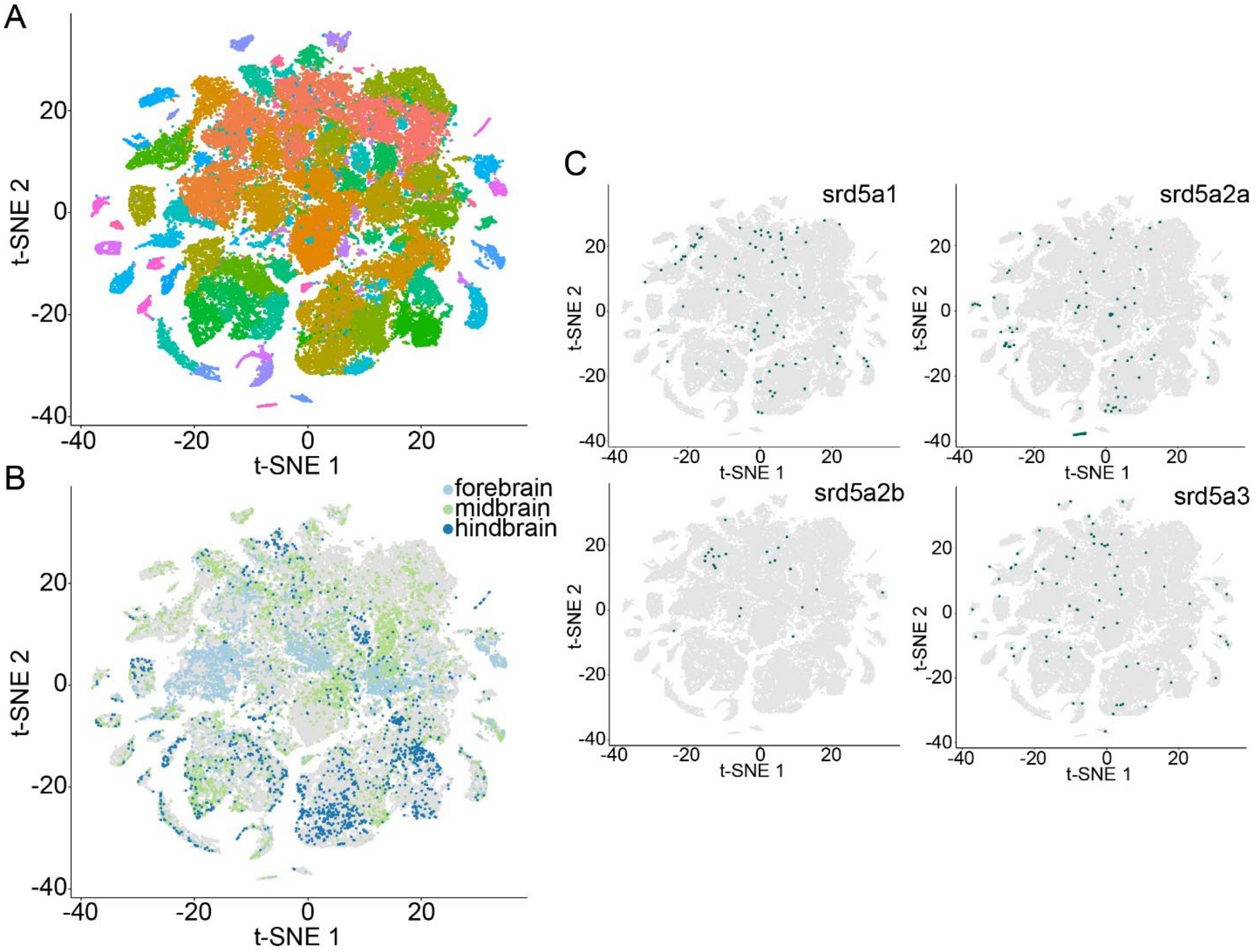


**Figure S2: Srd5a family members display scattered expression in multiple brain regions.**

**A**. t-SNE analysis plots of the single cell transcriptome of 23-25 days post fertilization zebrafish brains (n=65,718) colored by cluster. **B**. Samples originating from manually dissected fore-, mid- and hindbrains are highlighted. **C**. Samples expressing srd5a family members are highlighted.

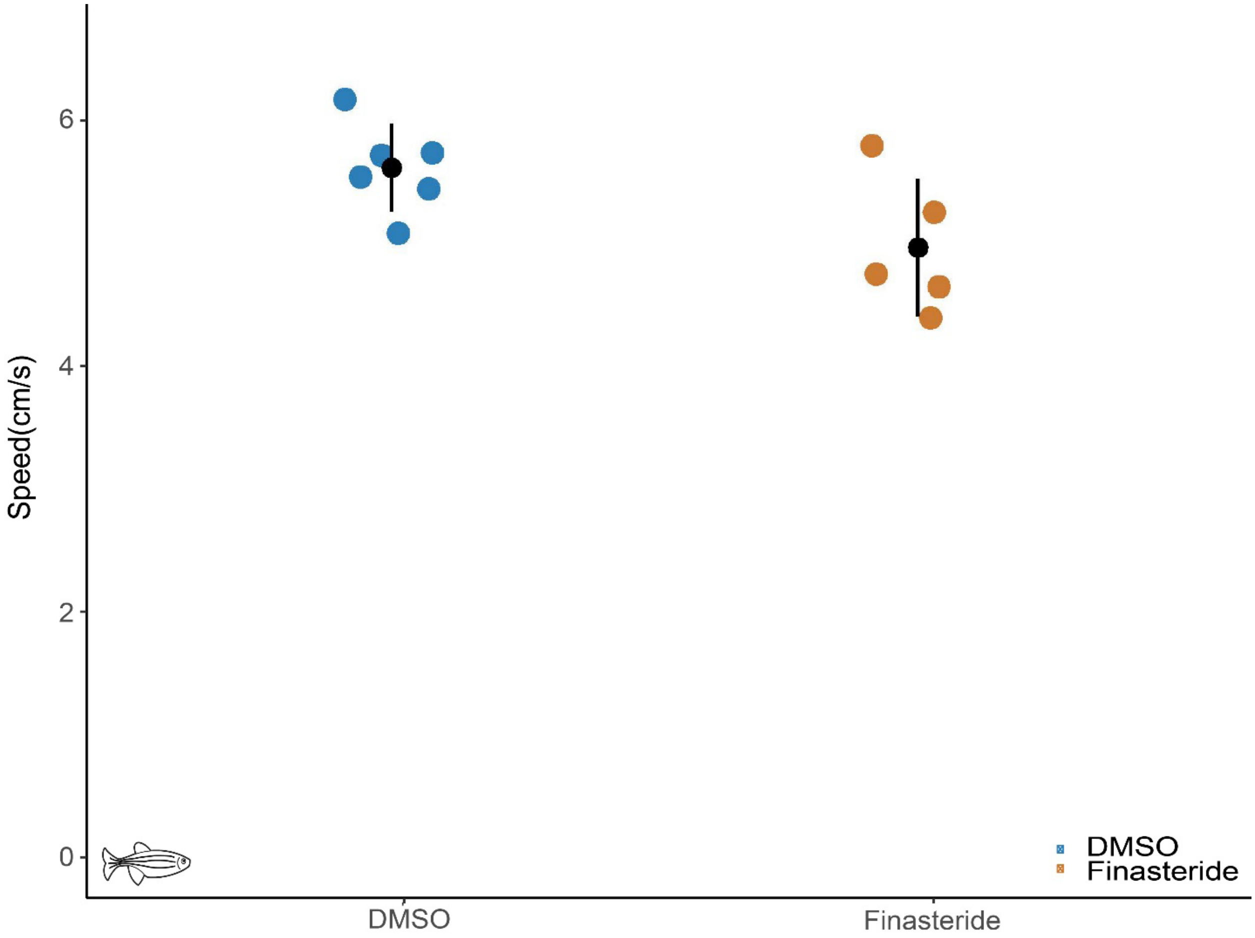


**Figure S3: Locomotion is unaffected in finasteride-treated fish.**

Data are the average speed of the animals in the arena (cm/s) from the different treatment conditions. DMSO n=6, Finasteride n=5. No significant difference. *p*-value computed by TukeyHSD on ANOVA (F(2,14)=2.77)p=0.1. Each n represents a group of 15 animals.

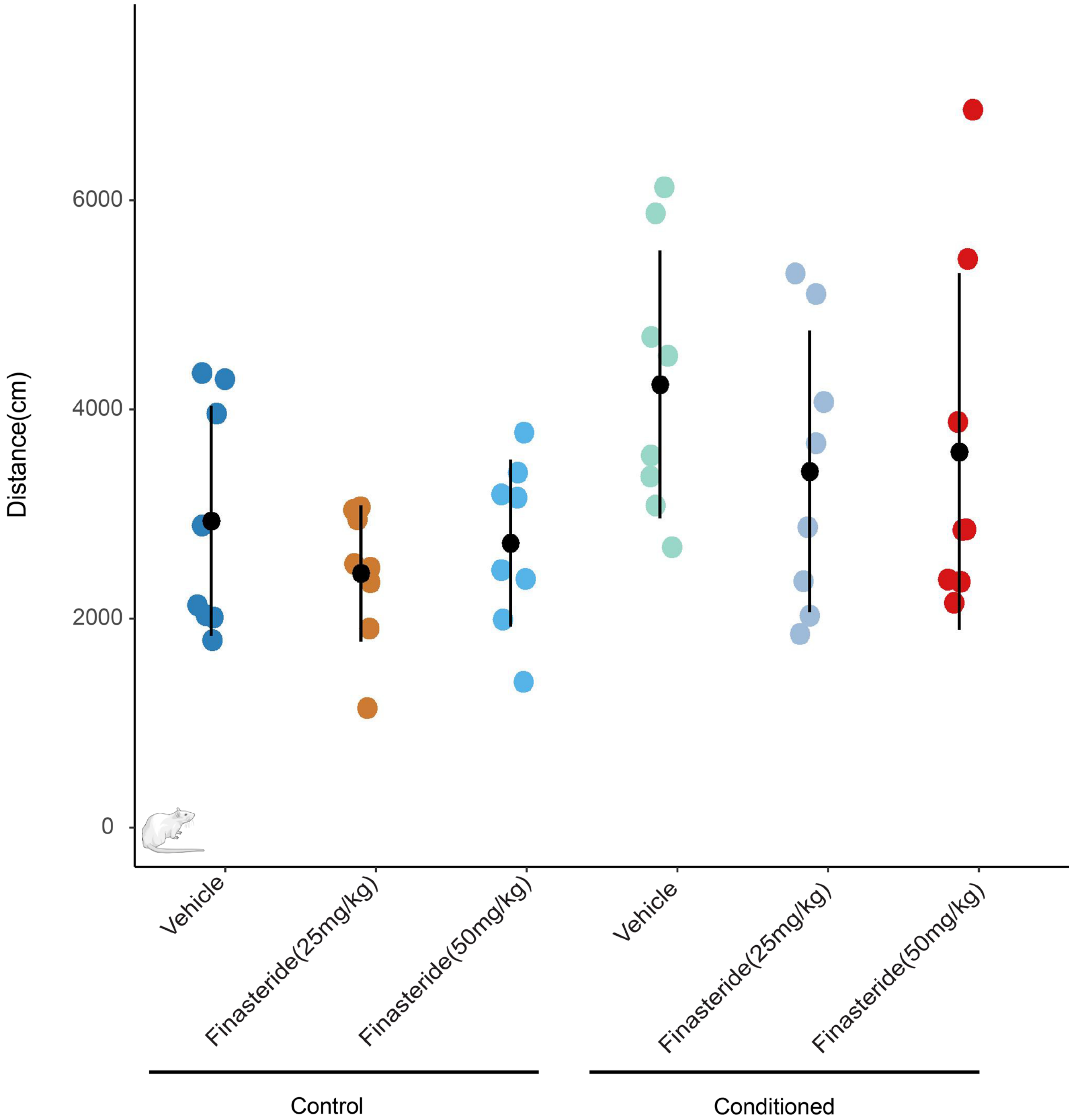


**Figure S4: Locomotion is unaffected in finasteride-treated rats**

Locomotion in the open-field test of rats treated with finasteride at 25 or 50 mg/kg, i.p.is not changed compared to corresponding vehicle-treated animals. N=8 per condition. Errors bars represent mean +/- s.e.m.

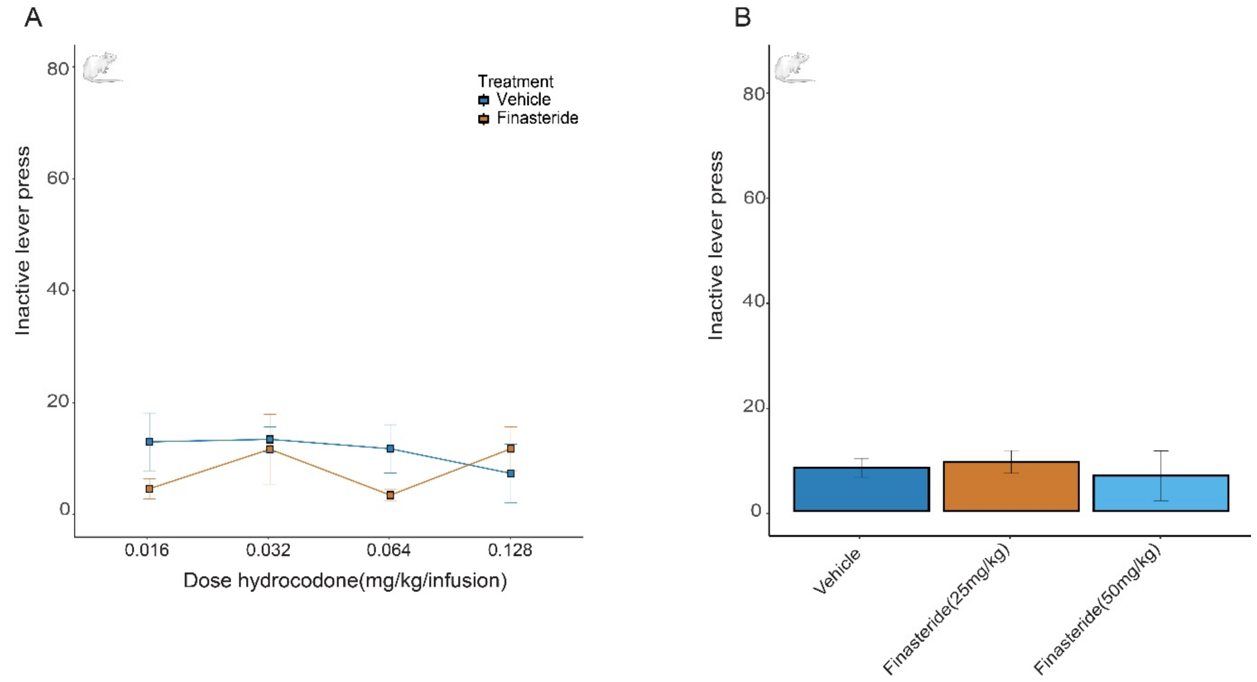


**Figure S5. No difference in inactive lever press during hydrocodone self-administration in rats A.** Pretreatment with 50mg/kg finasteride did not alter the number of inactive lever presses, regardless of the dose of hydrocodone. n=6 per condition Two-way ANOVA($F(3,38)=0.07, p=0.97$) **B.** Different doses of finasteride did not affect inactive lever presses for animals conditioned with 0.064mg/kg of hydrocodone. ANOVA($F(2,21)=0.19$)$p=0.82$. error bars represent mean +/- s.e.m.

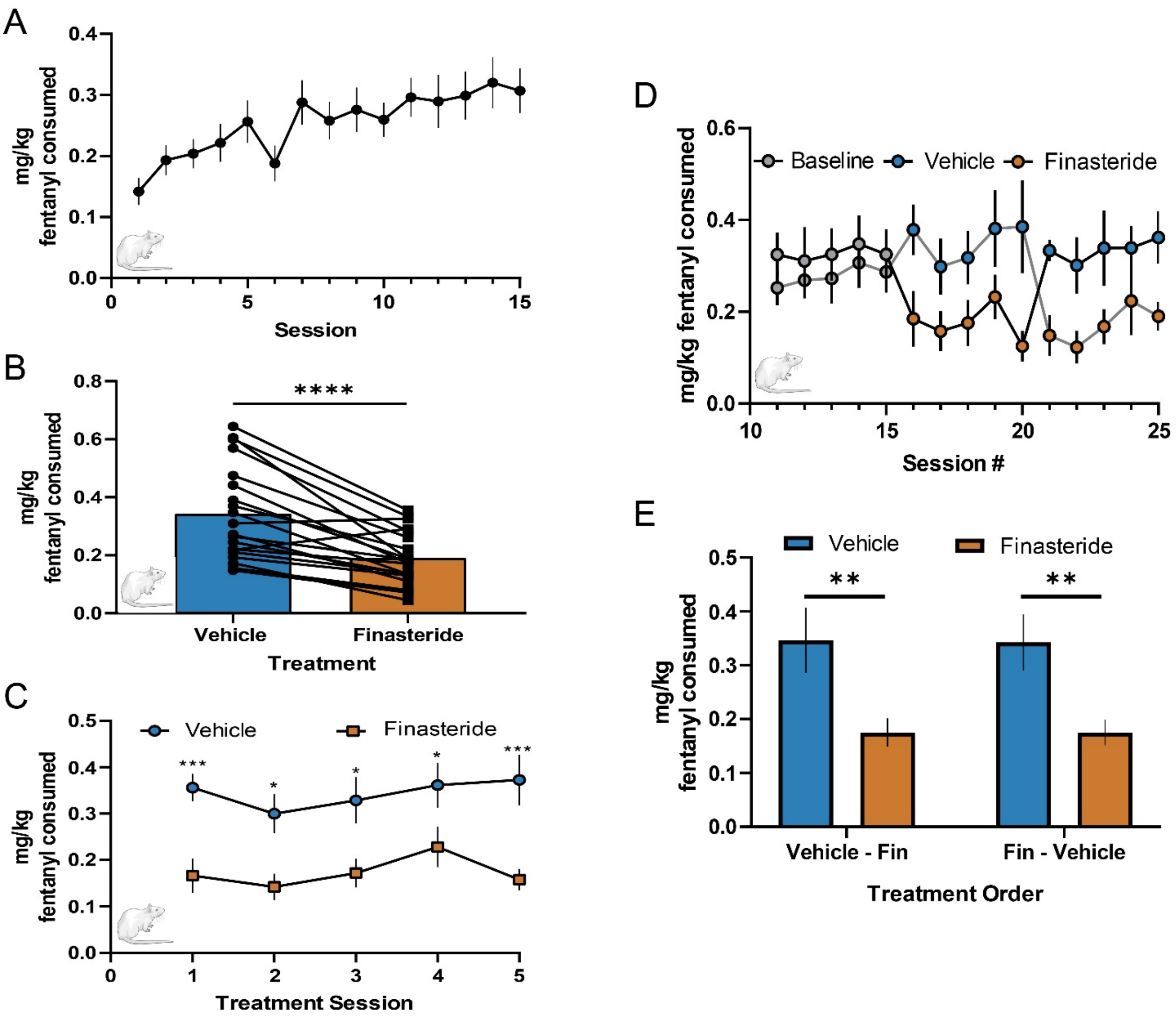


**Figure S6. Finasteride decreases fentanyl consumption in rats.**

**A.** Average fentanyl consumed in mg/kg for each baseline fentanyl self-administration session (n=20 rats). **B.** Average mg/kg of fentanyl consumed during vehicle (blue) and finasteride (orange) pretreated sessions. A paired t-test revealed finasteride treatment decreased the amount of fentanyl consumed compared to vehicle (t(19)=5.720)p<0.0001. **C.** Daily injections of finasteride (50mg/kg,IP; orange squares) significantly decreased fentanyl consumption compared to daily injections of vehicle (blue circles). A main effect of drug treatment was observed, Sidak post-hoc analysis performed for multiple comparisons on mixed-effect model (F(1,19)=26.94)p<0.0001. **D**

**& E.** Animals either received injections of finasteride during self-administration sessions 16-20 (n=10 rats, D: black line, E: Fin–Vehicle) or during sessions 21-25 (n=10 rats, D: gray line, E: Vehicle–Fin). **D.** Baseline refers to animals responding during sessions 10-15. **E.** The order of treatment had no effect on fentanyl consumed (F(1,18) = 0.0017, p=0.9670). No statistical interaction was observed between treatment and treatment order (F(1,18)=0.0025)p=0.9604. A main-effect of finasteride treatment was present (F(1,18)=22.85)p = 0.0001. P-values corrected for multiple comparisons on two-way ANOVA. *p*-value < 0.05, ****p*-value<0.001, *****p*-value<0.0001. Error bars represent the mean +/- s.e.m.

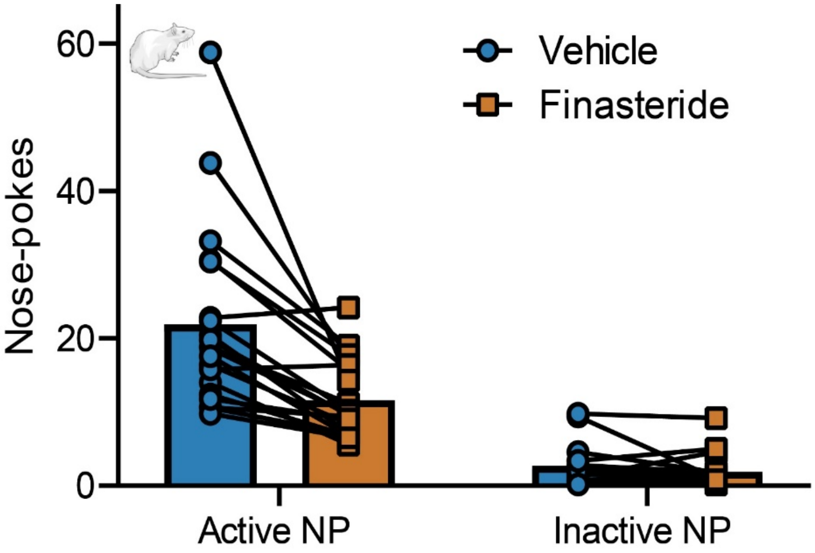


**Figure S7: No difference in inactive nose pokes in fentanyl-conditioned rats**

Finasteride didn't affect the number of nose pokes (NP) at the inactive port. Average operant nose-poke responses at both the active and inactive ports during finasteride (orange) and vehicle (blue) treatment sessions. N=20.

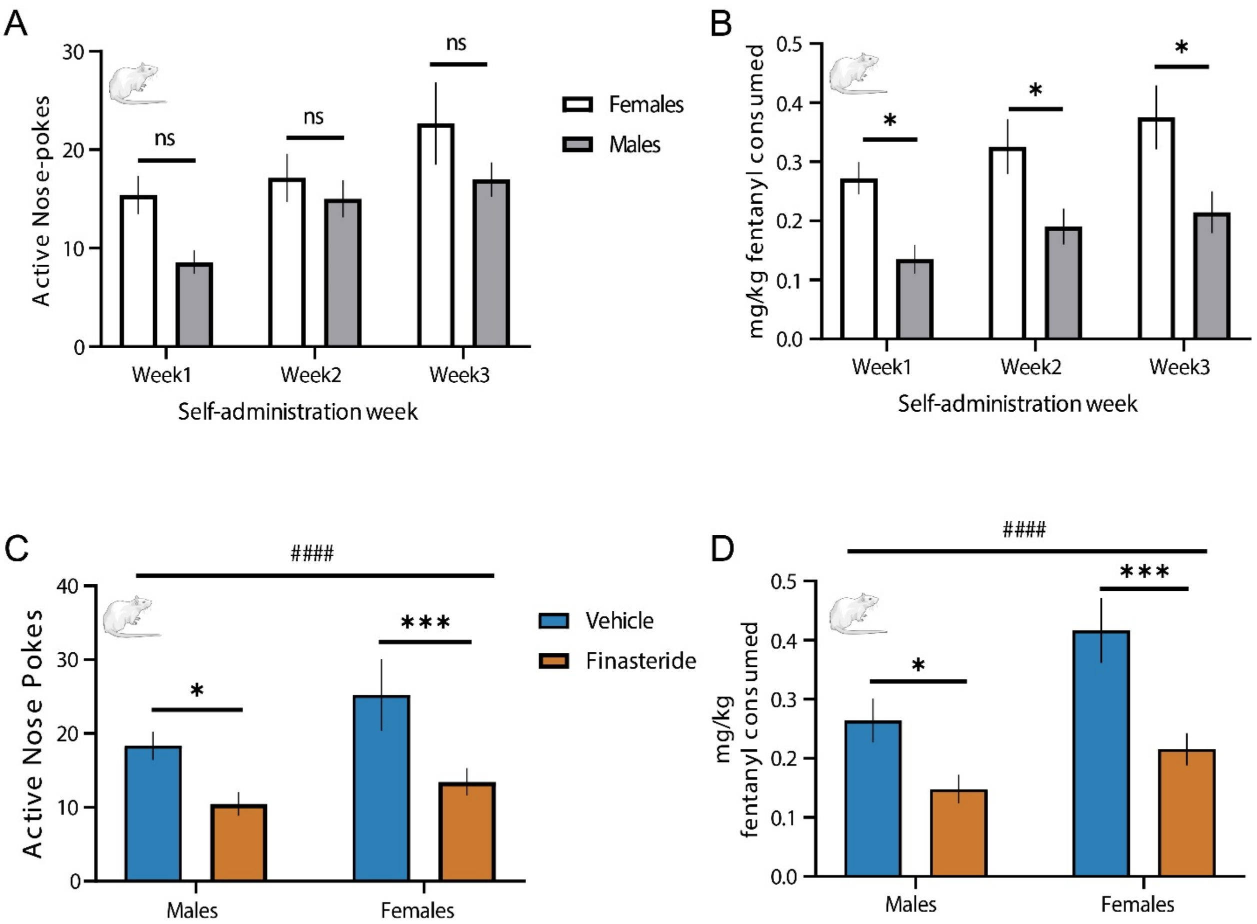


**Figure S8. Finasteride decreases oral fentanyl self-administration behaviors in male and female rats.**

**A.** Average operant nose-poke responses per session at the active port within each week of baseline fentanyl self-administration in male (filled bars) and female (open bars) Wistar rats. There was no observed statistical effect of sex on active nose-pokes (F(1,17)=2.821)p=0.1113 nor was there a significant self-admin week by sex interaction (F(2,34) = 0.9610)p=0.3927. A significant main effect of self-admin week was observed (F(2,34) = 9.848)p = 0.0004. **B.** Average mg/kg fentanyl consumption per session during each baseline week of oral fentanyl self-administration grouped by sex. There was no significant interaction between self-admin week and sex (F(2,34) = 0.1499). Statistical significance was observed in both main effects of sex (F(1,17) = 10.00)p = 0.0057 and self-admin week ($F(2,34) = 6.299$)$p = 0.0047$. A Sidak post-hoc analysis revealed female rats on average consumed more mg/kg of fentanyl in each session (?) during each week of baseline fentanyl self-administration. **C.** Average operant nose-poke responses during finasteride (orange) and vehicle (blue) treatment sessions. There was no difference in the effect of finasteride on active nose-pokes between the sexes, as evidenced by the lack of a significant sex x treatment interaction in the two-way ANOVA ($F(1,18) = 1.148$)$p= 0.2981$. An overall main effect of drug treatment was observed ($F(1,18) = 29.22$)$p < 0.0001$. **D.** Average mg/kg fentanyl consumption during finasteride (orange) and vehicle (blue) treatment sessions. There was no observed sex difference in the effect of finasteride on mg/kg of fentanyl consumed as no significant interaction was present between sex and treatment in a two-way ANOVA ($F(1,18)=2.069$, $p=0.1674$). There were main effects of both sex ($F(1,18)=6.110$, $p=0.0237$) and treatment ($F(1,18)=28.82$, $p<0.0001$). **p*-value < 0.05, ****p*-value<0.001, ##*p*-value<0.01 and indicates a main effect of treatment. Error bars represent the mean +/- s.e.m.

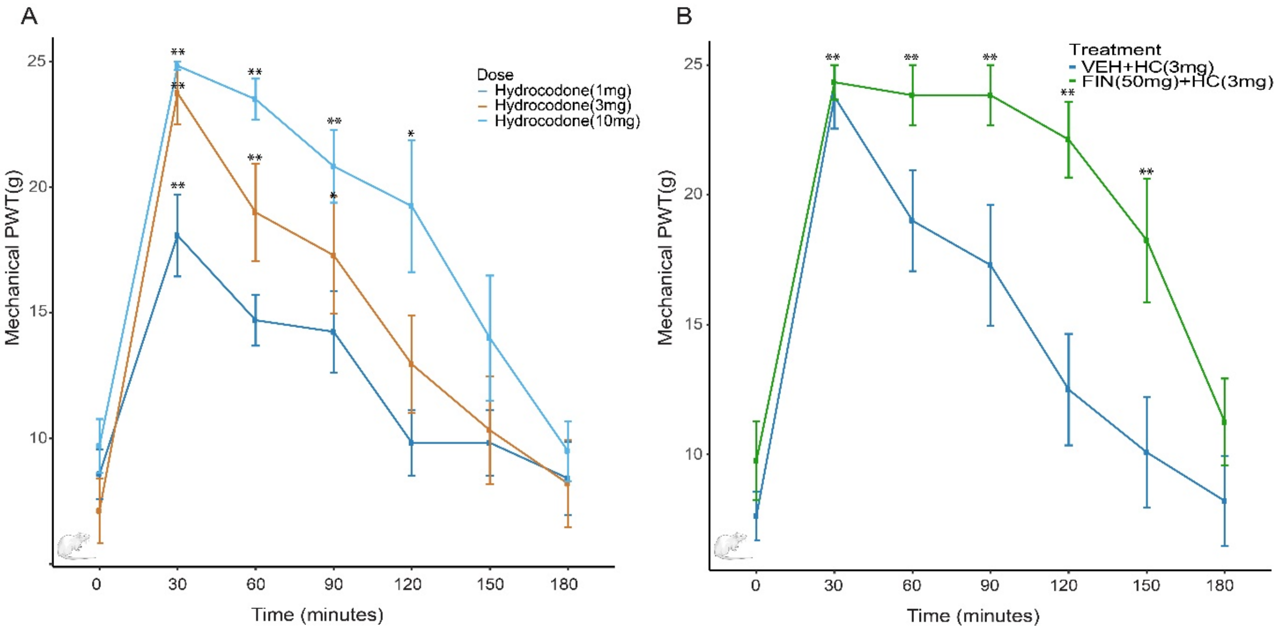


**Figure S9. Randall Sellito hydrocodone and finasteride.**

**A**. Paw withdrawal thresholds (PWT) Randall-Selitto assay. The withdrawal thresholds to paw pressure were measured up to 180 minutes after treatment with different doses of hydrocodone. Different doses of hydrocodone significantly increase latency to withdrawn the paw when a pressure in g is applied, compared to animals tested immediately after injection. Two-way ANOVA significant for TimeF(6,90)=53,90)p<0.0001 and Treatment F(2,15)=4.499)p=0,0295. N=6 per conditions **B**. Co-treatment with Finasteride (50mg/kg) did not block the antinociceptive effect of hydrocodone(10mg/kg) N=6 per condition. P compared with values measured before injection(Time 0)*p-value < 0.05, ** p-value<0.01.

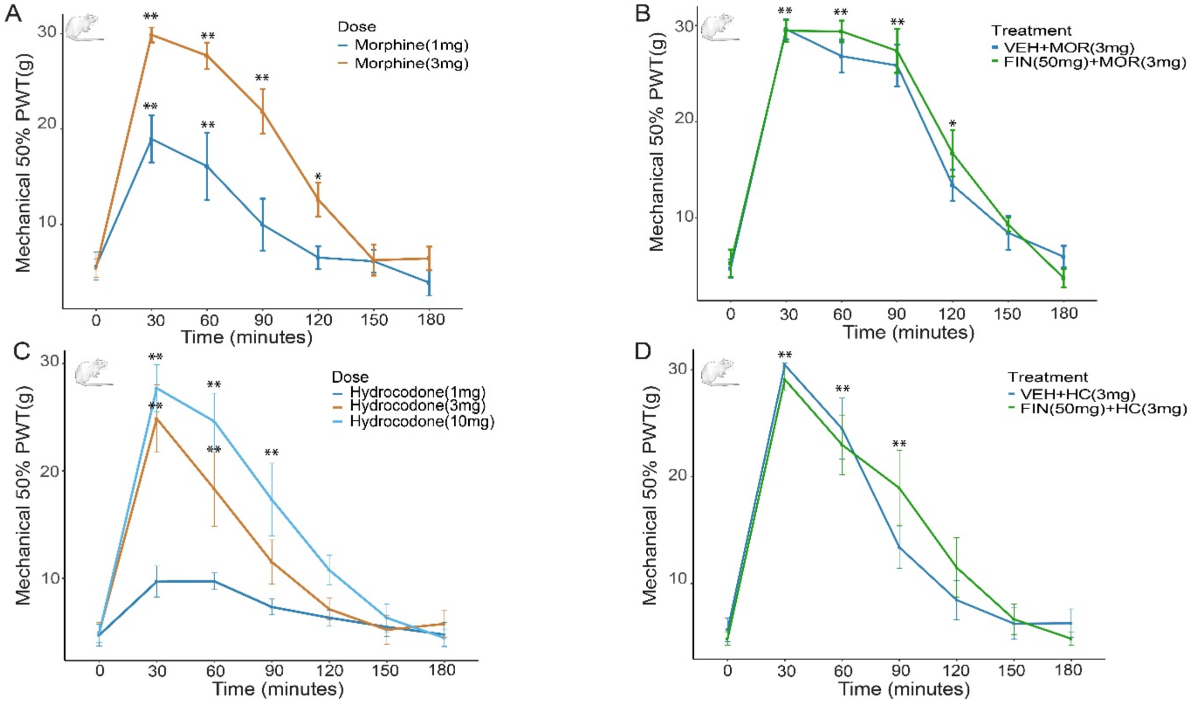


**Figure 10. Von-Frey morphine and hydrocodone and finasteride**

Finasteride does not affect the antinociceptive effect of opioids in a neuropathic pain model; paw withdrawal thresholds (PWT) to Von Frey filament. **A**. Different doses of morphine increase the PWT threshold. Two-way ANOVA revelaed a significant effect of time (F(6.72)=56.41)p<0.0001 and treatement (F(1.12)=14.40)p=0.0026. 1mg/kg n=6, 3mg/kg n=8 **B**. Co-treatment with finasteride(50mg/kg, IP) did not block the antinociceptive effect of morphine(3mg/kg). Two-way ANOVA significant effect of time (F(6.72)=125.4)p<0.0001, but no effect of treatment (F(1.12)=0.3659)p=0.5565 and no significant interaction (F(6,72)=0.4152)p=0.8666. Vehicle n=6, Finasteride n=8 **C**. Hydrocodone at doses of 3 and 10 mg/kg increased the PWT to Von Frey filament. N=6 per condition Two-way ANOVA identified significant main effects of time (F(6.126)=56.67)p<0.0001 and treatment (F(2,21)=9.488)p=0.0012. **D**. Co-injection with finasteride(50mg/kg) did not affect the antinociceptive activity of hydrocodone (10mg/kg), as the two-way ANOVA revealed a significant effect of time (F(6.72)=60.06)p<0.0001, but no main effect of treatment (F(1,12)=0.04560)p=0.8345 and no significant treatment x time interaction (F(6,72)=1.374)p=0.2368. Vehicle n=6, Finasteride n=8. P compared with values measured before injection(Time 0), **p*-value < 0.05, ** *p*-value<0.01.

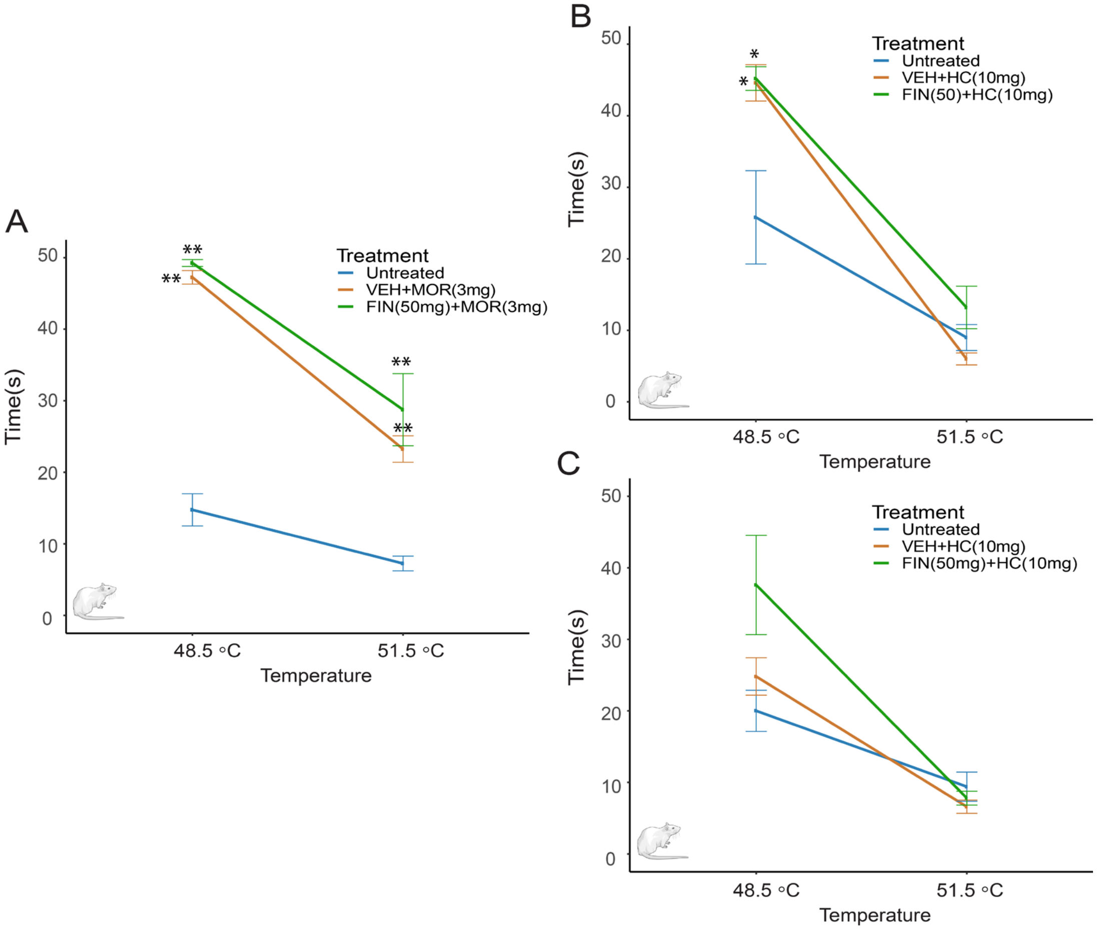


**Figure S11. Hot-plate morphine and hydrocodone and finasteride.**

Finasteride didn't affect the thermal antinociceptive effect of hydrocodone, as measured by the time before paw lick/retraction at different temperatures. *p* value compared with untreated animals. **A**. Finasteride didn't affect the thermal antinociceptive effect of morphine administered 60 min prior to the test, which measured the time before paw lick at different temperatures. 48.5°C :Two-Way ANOVA significant effect of treatment (F(2.9)=770.3)p<0,0001. 51.5°C two way-ANOVA significant effect of treatment (F(2.12)=7.66)p=0.0072. Both vehicle+morphine and finasteride+morphine are significantly different from untreated animals. No difference observed between vehicle+morphine and Finasteride-Morphine. **B**. Finasteride did not affect the thermal antinociceptive effect of hydrocodone administered 30 min prior to the test, which measured the time before paw lick at different temperatures. Two-Way ANOVA significance effect of treatment $(F(2.12)=6.1119)p=0.0147$ N=5 per condition. Both vehicle+hydrocodone and finasteride+hydrocodone are significantly different from untreated animals at 48.5°C. No difference observed between vehicle+hydrocodonde and finasteride+hydrocodone. **C**. Hydrocodone given 60 min prior to the test did not have a thermal antinociceptive. N=5**p*-value $< 0.05$, ** *p*-value$<0.01$.

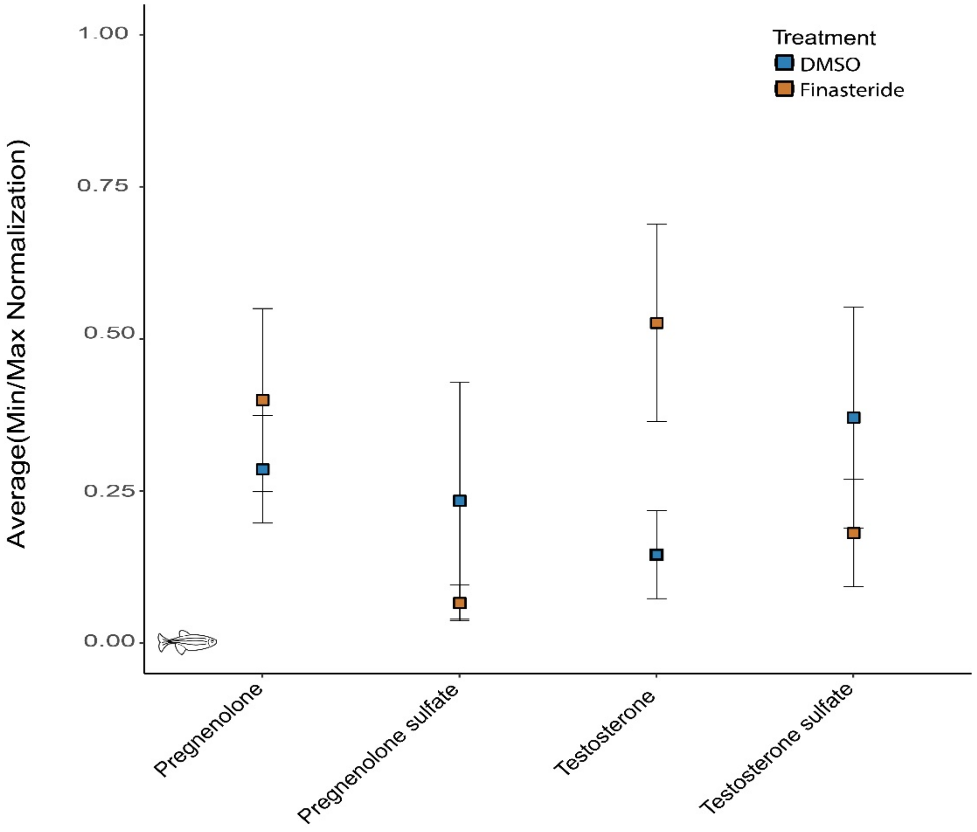


**Figure S12. Other sulfated steroids are not accumulating in finasteride-treated fish**.

Other sulfated steroids were not affected by the treatment with finasteride. Data are the normalization scores for the quantification of steroids in conditioned brains treated with DMSO or finasteride (10 µM). N=5 per condition. Each n represents a set of 10 brains.
